# Integrated scRNA-seq analysis identifies conserved transcriptomic features of mononuclear phagocytes in mouse and human atherosclerosis

**DOI:** 10.1101/2020.12.09.417535

**Authors:** Alma Zernecke, Florian Erhard, Tobias Weinberger, Christian Schulz, Klaus Ley, Antoine-Emmanuel Saliba, Clément Cochain

## Abstract

**Rationale:** Accumulation of mononuclear phagocytes (monocytes, macrophages and dendritic cells) in the vessel wall is a hallmark of atherosclerosis. Although single-cell RNA-sequencing (scRNA-seq) has shed new light on immune cell transcriptional diversity in atherosclerosis, it is still unknown whether the transcriptional states of mononuclear phagocytes are conserved between mouse and human atherosclerosis.

**Objective:** To integrate and compare macrophage and dendritic cell transcriptomes in mouse and human atherosclerosis.

**Methods and results:** We integrated 12 scRNA-seq datasets of immune cells isolated from healthy or atherosclerotic mouse aortas, and scRNA-seq data from 11 patients (n=4 coronary vessels, n=7 carotid endarterectomy specimens) from two independent studies. Integration of mouse data recovered previously described macrophage populations and identified novel subpopulations with discrete transcriptomic signatures within populations of aortic resident (*Lyve1*), inflammatory (*Il1b*), as well as foamy (*Trem2^hi^*) macrophages. We identified unique transcriptomic features distinguishing aortic intimal resident macrophages from atherosclerosis-associated *Trem2^hi^* macrophages. Also, populations of *Xcr1^+^* type 1 classical dendritic cells (cDC1), *Cd209a^+^* cDC2 and mature DCs (*Ccr7, Fscn1*) were detected. In humans, we uncovered macrophage and dendritic cell populations with gene expression patterns similar to those observed in mice in both vascular beds. In particular, core transcripts of the *foamy/Trem2^hi^* signature (*TREM2, SPP1, GPNMB, CD9*) mapped to a specific population of macrophages in human lesions. Cross-species data integration demonstrated transcriptionally proximal macrophage and dendritic cell populations in mice and humans.

**Conclusions:** We demonstrate conserved transcriptomics features of macrophages and dendritic cells in atherosclerosis in mice and humans, emphasizing the relevance of mouse models to study mononuclear phagocytes in atherosclerosis.

## Introduction

Atherosclerosis is a chronic disease of the arterial wall characterized by chronic lipid accumulation and inflammation in the vascular intima, and its main clinical manifestations, myocardial infarction and ischemic stroke, together constitute the most frequent cause of death worldwide (1). Many adaptive and innate immune cell types have been proposed to contribute to vascular inflammation and atherosclerosis (2), and accumulation of macrophages and lipid-laden macrophage foam cells are hallmarks of atherosclerotic lesions (3). As lesional macrophages perform both atheroprotective (e.g. efferocytosis, lipid clearance) and atherogenic (inflammatory cytokine secretion, proteolysis) functions, it has long been assumed that functionally distinct macrophage subsets populate atherosclerotic vessels. Recent studies employing single-cell RNA-sequencing (scRNA-seq) analyses of the vascular immune cell infiltrate have shed new light on macrophage diversity in atherosclerosis (4), (5).

In mice, several independent scRNA-seq studies have evaluated the transcriptomic identity of aortic macrophages in experimental models of atherosclerosis (6–11), and identified heterogeneous disease-associated macrophage populations, with non-foamy macrophages characterized by a pro-inflammatory gene expression profile (Inflammatory macrophages), and lipid-laden foamy macrophages showing low expression of inflammatory genes, high expression of the myeloid receptor *Trem2* and of a set of genes involved in lipid metabolism or lysosome function (foamy/*Trem2^hi^* macrophages)(10, 11). We in addition have described macrophages with features of adventitial resident cells (*Lyve1* expression (12, 13)) in control and atherosclerotic aortas (Resident/Resident-like macrophages) (6). Macrophages with a strong type I interferon response signature, similar to interferon inducible cells (IFNICs) found in the ischemic heart (14), were also observed in atherosclerotic aortas (7), (9). A recent study, combining scRNA-seq analyses, fate mapping and imaging experiments, furthermore demonstrated the existence of a self-renewing aortic intimal resident macrophage (Mac-AIR) population in the normal mouse aorta, characterized by the expression of CD11c, MHCII and a specific transcriptomic signature (15). First available data in humans suggest that macrophages with distinct transcriptional states are also found in atherosclerotic vessels (16), (17), (18).

As mouse models of atherosclerosis are widely employed to decipher the pathogenesis of atherosclerosis, and to perform pre-clinical investigations of new potential therapeutic targets (19), it is critical to determine whether macrophage transcriptional states in atherosclerotic vessels are conserved across mouse models of atherosclerosis and whether the transcriptional states of macrophages in mouse atherosclerosis are of relevance to human disease. Recently developed computational tools allow integration of scRNA-seq data across independent studies, technological platforms, and species (20, 21), thus providing an unprecedented opportunity to compare cellular states across experimental conditions, and from animal disease models to cells directly obtained from diseased tissue from patients (22).

Here, we performed a computational integration of scRNA-seq data of immune cells from independent studies investigating mouse models of atherosclerosis and human atherosclerotic plaque tissue, using the data integration features of the Seurat v3 package (20). Our work shows conserved transcriptomic features of macrophages in atherosclerosis across mouse models and identifies novel putative atherosclerosis-associated macrophage subpopulations. We also observed conserved dendritic cell populations in mouse and human atherosclerotic tissue. We further provide evidence that major features of macrophage states observed in mouse atherosclerosis are conserved in human lesions, emphasizing the relevance of mouse models to study macrophage biology in atherosclerosis.

## Methods

Using 12 distinct mouse scRNA-seq datasets from 6 studies (6–9, 23), (15) (**Supplementary Table 1**), and 3 human scRNA-seq datasets from two studies (17, 24), we performed three integrated analyses (**Figure SIA-C**): (i) integration of all mouse data, (ii) integration of human mononuclear phagocyte data, and (iii) integration of mouse and human mononuclear phagocyte data. Deposited cell-gene count matrixes were analyzed in Seurat (20), where all datasets were first pre-processed individually for quality control filtering, selection of relevant cells for further analysis, and assignment of metadata information (e.g. species, protocol, patient) (**Supplementary Table 1**, **Figure SI**). To identify the sex of mice used in some studies where this information was unavailable (9), we interrogated the expression of X chromosome (*Xist*) or Y chromosome specific transcripts (*Ddx3y, Uty, Eif2s3y*) (**Figure SID**). Most studies employed male mice, except *Apoe^-/-^* studies by Winkels et al. that used females (8), while Lin et al. (9) employed male mice for the atheroprogression and female mice for the atheroregression studies (**Figure SID**, **Supplementary Table 1**). Integration was performed using Seurat v3 (20), essentially according to the “Standard Workflow” protocol provided by the authors (https://satijalab.org/seurat/v3.1/integration.html). The code used for analysis will be provided as R notebooks. We have mostly limited data interpretation to qualitative aspects of cell transcriptional states and did not interpret quantitative changes (e.g. proportion of certain types of macrophages) beyond their presence/absence across experimental conditions or species. This is based on limitations of the individual studies, including lack of experimental replicates (mouse studies were performed with n=1 scRNA-seq run per condition), low number of patients and very low number of cells analyzed in some patients, as well as the poorly controllable differences in cell type recovery by tissue digestion and cell processing methods between studies. Overlapping marker gene lists shared between human macrophage clusters and their putative mouse counterparts were identified using InteractiVenn (25).

## Results

### Integrated analysis of mouse aortic immune cells scRNA-seq datasets

The integrated analysis of mouse aortic leukocyte datasets from different models of atherosclerosis and experimental conditions (**Table 1**, **Figure SI**) allowed us to analyze a final number of 22,852 cells (**Figure 1A, B**). We identified macrophages and dendritic cells based on the expression of canonical markers such as *Adgre1* (encoding F4/80), *Fcgr1* (encoding CD64) and *Itgax* (encoding CD11c) (**Figure 1C**). Several populations of T (*Cd3d*), B (*Cd79a*), Cytotoxic/NK cells (*Nkg7*) and neutrophils (*S100a8, S100a9*) could be discerned. A cluster resembling type 2 Innate Lymphoid Cells (ILC2) (*Il1rl1, Gata3*) was identified, possibly also containing mast cells and basophils (*Cpa3, Calca*) (**Figure SIIA**). We had previously identified these cells as mixed/mast cells in (6). A cluster of non-leukocytic cells was also found, originating from the scRNA-seq analysis of intimal BODIPY^+^ foam cells, which included lipid rich intimal cells of both immune and non-immune origin (7).

**Figure 1:**
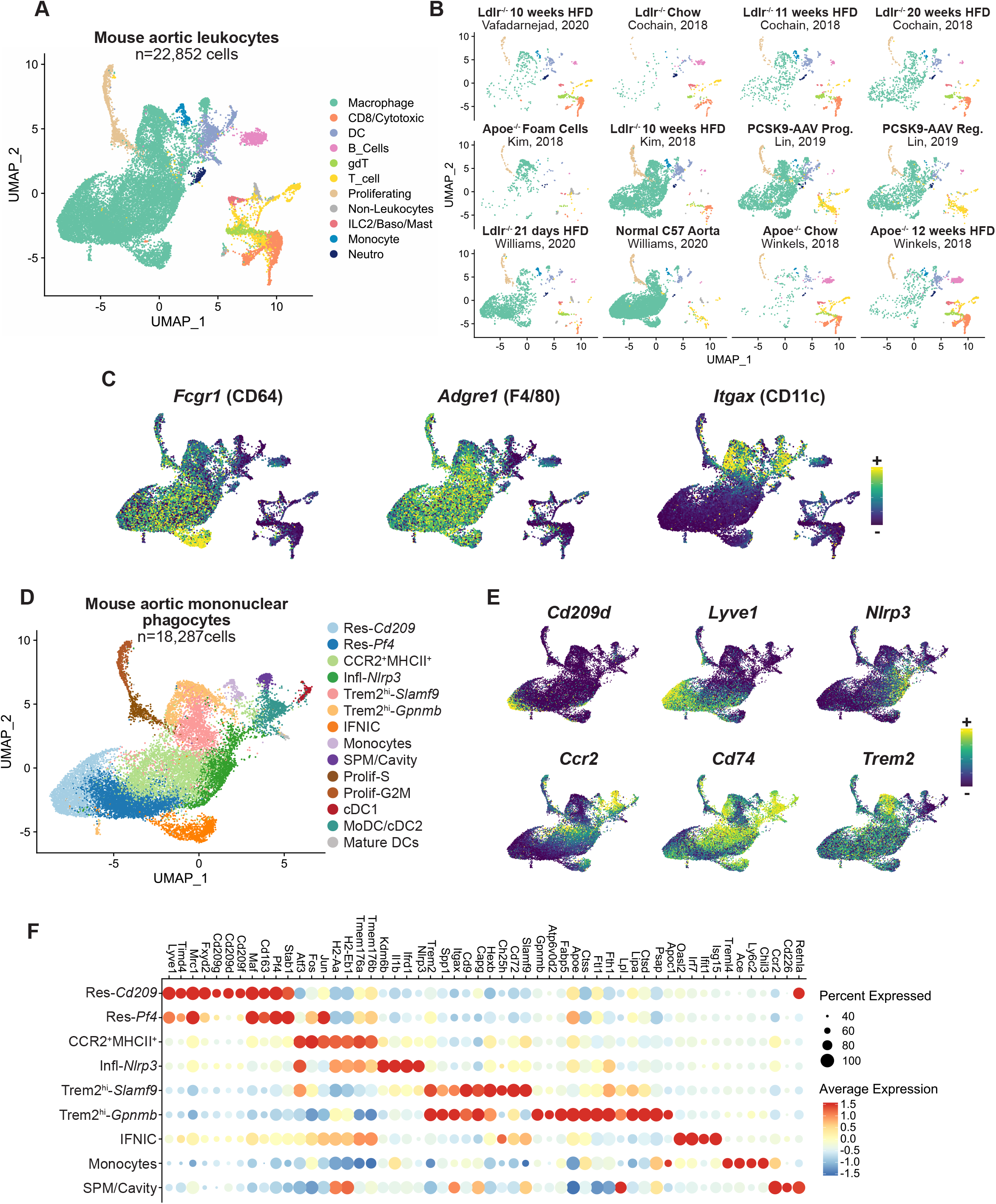
integrated scRNA-seq analysis of vascular inflammation in mouse atherosclerotic aortas. **A)** UMAP representation of integrated scRNA-seq gene expression data in 22,852 cells from mouse atherosclerotic aortas with identification of the major immune cell lineages (DC: dendritic cells; gdT: gammadelta T cells); **B)** projection of single cells in the UMAP space according to dataset and experimental condition of origin. **C)** Expression of the indicated transcripts projected onto the UMAP plot. **D)** UMAP plot of the mononuclear phagocyte data subset with clustering analysis; **E)** expression of the indicated transcripts projected onto the UMAP plot; **F)** dot plot of average gene expression of the indicated marker transcripts in the macrophage clusters.

### Subpopulations of Resident/Resident-like and Inflammatory macrophages in atherosclerotic aortas

Focusing our analysis on mononuclear phagocytes, we then performed clustering of cells corresponding to monocytes, macrophages, dendritic cells and proliferating cells at higher resolution to identify potential sub-populations within macrophages and dendritic cells (**Figure 1D**). Resident/Resident-like macrophages (*Lyve1, Timd4, Mrc1, Pf4*) (**Figure 1D-F**) could be divided into two clusters, one of which showed higher expression of *Cd209d*, *Cd209f* and *Cd209g* (Res-Cd209) (**Figure 1D-F**), corresponding to Cd209^+^ resident macrophages identified by Cole et al. (26). Macrophages corresponding to previously described ‘Inflammatory Mφ’ also comprised two clusters: Inflammatory-*Nlrp3* macrophages displayed high *Nlrp3, Il1b* or *Kdm6b* expression, while a second subset of CCR2^+^MHCII^+^ macrophages expressed *Ccr2*, MHCII encoding transcripts (*Cd74, H2-Aa, H2-Eb1*) and was enriched for *Tmem176a* and *Tmem176b* (**Figure 1D-F**). Proliferating cells could be delineated into S-phase and G2M-phase cells by applying cell cycle scoring in Seurat (20) (**Figure 1D, Figure SIIB**). In addition to major macrophage subsets, we observed populations consistent with a previous integrative analysis (5), including cells with a gene expression profile characteristic of small peritoneal macrophages (SPM/Cavity cluster: *Itgax^+^Cd226^+^Ccr2^+^MHCII^+^*) (5), type I interferon response cells (IFNIC cluster: *Isg15, Oasl2*), and monocytes expressing genes characteristic of Ly6C^hi^ (*Ly6c2, Chil3, Ccr2*) and Ly6C^low^ (*Ace*, *Treml4*) subsets (28) (**Figure 1D and F**). Immediate early genes (IEGs) expression can be induced in mononuclear phagocytes during tissue digestion and processing (27). Based on the expression of 18 IEGs, we applied an IEG expression score to macrophages and dendritic cells, and cells with the highest score mapped to the ‘inflammatory Mφ’ clusters (**Figure SII C**). However, stress-induced gene expression is unlikely to have caused a major bias in our analysis, as cell clustering was not substantially affected by regressing out variability caused by expression of immediate early genes (**Figure SII D-F**).

### Subpopulations of Trem2^hi^ macrophages and aortic intimal resident macrophages

We identified two clusters enriched for genes characteristic of the *foamy/Trem2^hi^* signature (10), (5): *Trem2, Spp1, Cd9, Itgax* (**Figure 1D-F**). The first population was further enriched for transcripts such as *Slamf9, Ch25h* and *Cd72 (Trem2^hi^-Slamf9*, **Figure 1D and F**). The second population (*Trem2^hi^-Gpnmb*) was enriched for *Gpnmb, Atp6v0d2* and transcripts characteristic of a foamy signature and TREM2-reponse genes (29), (30) (*Lpl*, *Lipa, Fabp5, Apoc1, Apoe*) (**Figure 1D and F**). Consistent with previous analyses (6),(7) and recent observations in *Apoe^-/-^Cx3cr1^GFP^Cd11c^YFP^* mice (31), foamy/Trem2^hi^ macrophages were enriched for *Itgax* (CD11c) (**Figure 1C and F**). Most intimal BODIPY^+^ foam cells of the immune lineage from *Apoe^-/-^* mice (7) mapped to UMAP coordinates corresponding to foamy/Trem2^hi^ macrophage clusters (**Figure 1B**).

Aortic intimal resident macrophages (Mac-AIR) have recently been described to populate the vascular intimal niche in the normal aorta and present a specific transcriptomic signature (15). Monocytes infiltrating the intima under atherogenic conditions have furthermore been proposed to acquire transcriptomic features resembling those of Mac-AIR, and to further upregulate lipid-metabolism related and TREM2-response associated transcripts (15), (30). To define the relationship of Mac-AIR with both *Trem2^hi^* populations, we first identified Mac-AIR in scRNA-seq data from healthy C57BL/6 aortas (15), and examined their position on the integrated data UMAP (**Figure 2A**). Mac-AIR from healthy aortas mapped mostly to the *Trem2-Gpnmb* (79 out of 107 cells, i.e. 73.8%) and the *Trem2-Slamf9* (23/107 cells, 21.5%) clusters (**Figure 2A**), in line with previous observations that Mac-AIR and intimal foamy macrophages are transcriptionally proximal (15). Mac-AIR mapped to an area of the UMAP characterized by high expression of MHCII-encoding transcripts (e.g. *Cd74, H2-Ab1, H2-Eb1* **Figure 1E**), consistent with MHCII expression by Mac-AIR (15). When clustering the integrated mononuclear phagocyte data using a higher resolution parameter in Seurat (20), we could identify an independent cluster mapping to Mac-AIR coordinates (‘Mac-AIR Signature’ cluster, **Figure 2B**). We analyzed differentially expressed genes between this ‘Mac-AIR Signature’ cluster, the *Trem2^hi^-Gpnmb* cluster, and the *Trem2^hi^-Slamf9* cluster. Expectedly, Mac-AIR where enriched for MHCII encoding transcripts, but also for other genes e.g. *Vcam1* and *Hes1* (**Figure 2C**). Relative to Mac-AIR and *Trem2^hi^-Gpnmb*, the *Trem2^hi^-Slamf9* cluster was enriched for *Nes*, *Cd72, Ch25h*, and inflammatory markers (*Tnfsf9, Il1b*) (**Figure 2C**), although these were expressed at lower levels than in bona fide pro-inflammatory macrophages (**Figure 1F**, and not shown). *Trem2^hi^-Gpnmb* had a specific signature including e.g. *Il7r, Psap* and *Fabp5* (**Figure 2C**). Importantly, Mac-AIR expressed lower levels of *Trem2, Spp1* and *Cd9* (**Figure 2D**), transcripts generally associated with the disease-associated “*Trem2^hi^*” signature in atherosclerosis (5),(6) and other diseases (30). Mac-AIR and *Trem2^hi^-Gpnmb*, but not *Trem2^hi^-Slamf9*, expressed *Acp5* (**Figure 2D**). A similar signature was obtained when comparing only Mac-AIR from healthy C57 aortas (15) versus the *Trem2^hi^-Gpnmb* and *Trem2^hi^-Slamf9* populations (not shown).

**Figure 2:**
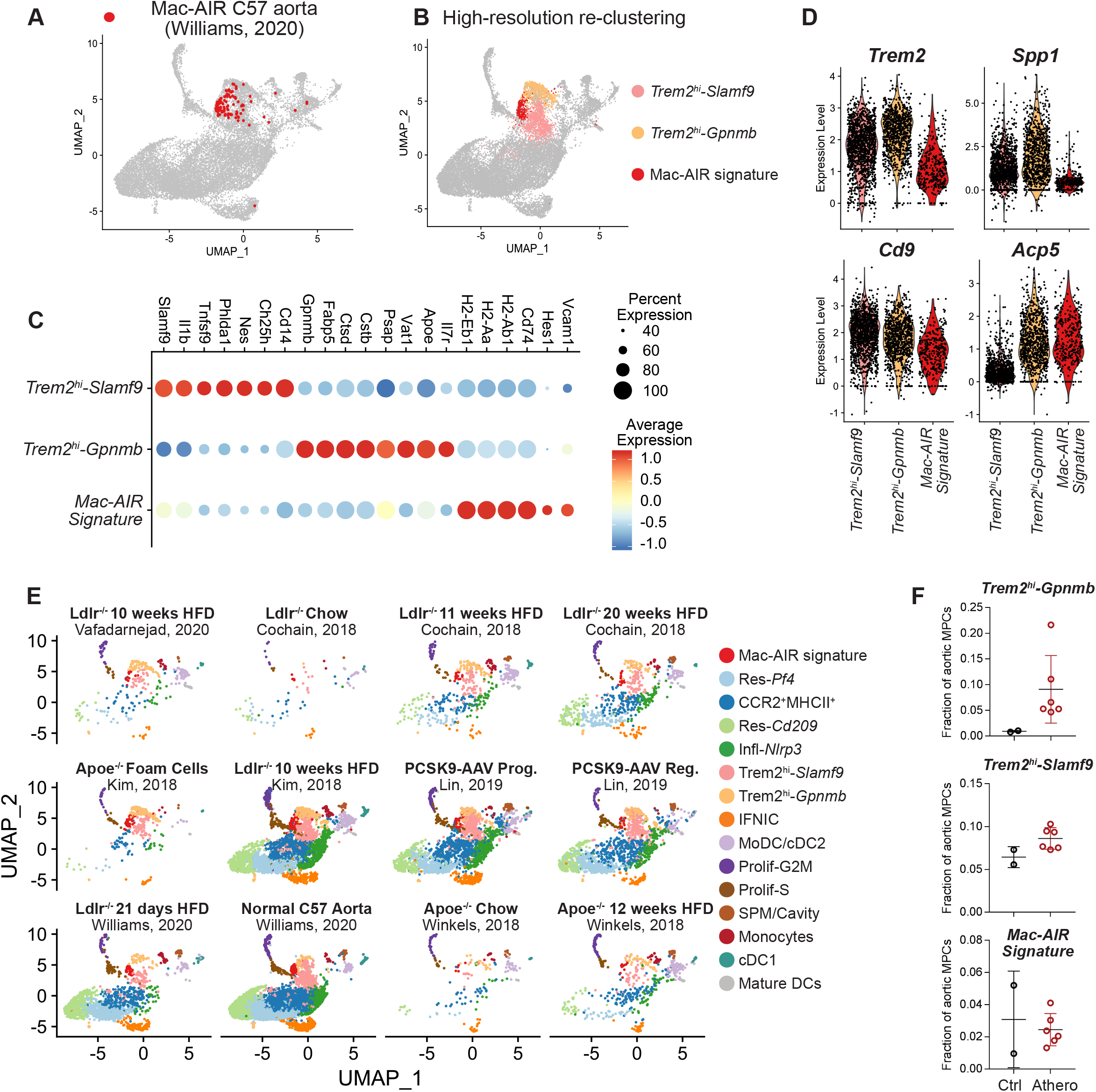
Characterisation of subpopulations within *Trem2^hi^* macrophages and their relationships to Mac-AIR. **A)** projection of cells corresponding to steady state C57BL6 aorta Mac-AIR cells (Williams, Nat Immunol 2020) on the UMAP plot and **B)** high-resolution re-clustering identifying an independent cluster with a Mac-AIR signature; **C)** dot plot showing average expression of selected marker genes in Mac-AIR, *Foamy/Trem2^hi^Gpnmb* and *Foamy/Trem2^hi^Slamf9* populations; **D)** expression of the indicated transcripts in Mac-AIR, Foamy/*Trem2^hi^Gpnmb* and Foamy/*Trem2^hi^Slamf9* populations; **E)** UMAP plot with identification of macrophage subclusters including the Mac-AIR signature cluster, split according to dataset of origin; **F)** fraction of Mac-AIR, Foamy/*Trem2^hi^Gpnmb* and Foamy/*Trem2^hi^-Slamf9* populations among total aortic mononuclear phagocytes (MPCs). For consistency, not all datasets were used in **F**: datasets from *Apoe^-/-^* mice (Winkels et al. 2018) were excluded as chow fed *Apoe^-/-^* mice do not represent real non-atherosclerotic controls; the ‘Kim Foam Cell’ dataset was excluded as it is technically enriched for foam cells; the 21 days HFD dataset from Williams et al. Nat Immunol 2020 was excluded as it represents a very early time point of lesion formation.

We then analyzed the presence of *Trem2^hi^-Gpnmb* and *Trem2^hi^-Slamf9* and Mac-AIR clusters across datasets (**Figure 2E**) and calculated their relative abundance among total mononuclear phagocytes in control and atherosclerotic aortas (**Figure 2F**, not all datasets were considered, see Figure 2 legend). *Trem2^hi^-Gpnmb* macrophages represented 1% or less of all mononuclear phagocytes in control aortas, and 4.6% to 21.6% in atherosclerotic arteries. *Trem2^hi^-Slamf9* cells represented 5-10% of all mononuclear phagocytes both in control and atherogenic conditions. Mac-AIR represented less than 6% of all aortic mononuclear phagocytes in all the conditions (**Figure 2F**). While these analyses are statistically limited given the low number of replicates and low cell numbers in some datasets, they nevertheless clearly confirm that *Trem2^hi^-Gpnmb* macrophages with a foamy macrophage gene expression signature are induced by atherogenic conditions.

*Trem2^hi^-Gpnmb, Trem2^hi^-Slamf9*, and Mac-AIR were predominant in the intimal foam cell dataset from Kim et al. (7) in the integrated analysis (**Figure 2E**), representing 38%, 14% and 9% of mononuclear phagocytes, respectively. Analysis of this dataset alone recovered three macrophage populations with expression signatures reminiscent of Mac-AIR (*Acp5, Cd74, Mmp12, Gngt2*), *Trem2^hi^-Gpnmb (Gpnmb, Fabp5, Cstb, Psap*), and *Trem2^hi^-Slamf9* (*Slamf9, Cd72, Cd14, Ch25h*) macrophages (**Figure SIIIA-B**), further indicating that *Trem2^hi^-Gpnmb,Trem2^hi^-Slamf9* and Mac-AIR contribute to intimal foamy macrophages.

Altogether, this integrated analysis shows that atherosclerosis-associated macrophage transcriptional states are conserved across experimental mouse models of atherosclerosis, and that discrete subpopulations exist within the main aortic macrophage subtypes. We further confirm that Mac-AIR that reside in the normal mouse intima share transcriptional similarities with *Foamy/Trem2^hi^* macrophages but express higher levels of transcripts encoding MHCII, express lower levels of *Trem2*, and do not express a specific set of transcripts characteristic of the disease-associated Foamy/*Trem2^hi^* signature.

### Aortic macrophage subsets in angiotensin-II induced vascular inflammation

To investigate whether similar macrophage states could be observed in a distinct context of aortic inflammation, we integrated the mononuclear phagocyte data (**Figure 1D**) from atherosclerosis studies with recently generated data in a model of angiotensin II-induced aortic inflammation also characterized by extensive macrophage infiltration and activation (32). All the mononuclear phagocyte population observed in atherosclerotic aortas were recovered in the aortic adventitia in the Angiotensin-II inflammation model (**Figure SIVA-C**). While pro-inflammatory CCR2^+^MHCII^+^ macrophages (also enriched for *Tmem176a/Tmem176b*, **Figure SIVC**)) dominated in this model (726 out of 2932 cells, 24.76%), we observed a prominent population of *Trem2^hi^-Gpnmb* cells (196 cells, 6.68%), indicating that these cells are also present during aortic inflammation in hyperlipidemia-independent models (**Figure SIVA-C**).

### Integrated analysis of aortic dendritic cells in mouse atherosclerosis

We identified cells corresponding to monocyte-derived DCs and/or cDC2 (MoDC/cDC2: *Cd209a, Clec10a, Ifitm1, Napsa*), cDC1 (*Xcr1^+^Clec9a^+^*), and *Fscn1^+^Ccr7^+^* mature DCs (**Figure 1D, Figure SVA**). We had previously identified the MoDC/cDC2 cluster as potential monocyte-derived dendritic cells based on the expression of *Cd209a* (33), and intermediate expression of monocytic markers (*Ccr2, Csf1ŕ*) (6). However, a recent report indicates that in inflammatory conditions, cDC2 can acquire an “inflammatory-cDC2” state with surface CD64 expression, that can be discriminated from monocyte-derived cells by expression of CD26 (encoded by *Dpp4*) and absence of expression of CD88 (encoded by *C5ar1*) (34). Recent work further suggests that CD88 can aptly discriminate monocyte/macrophages from dendritic cells in mice and humans (35). In atherosclerotic aortas, the MoDC/cDC2 cluster expressed *Dpp4* (CD26) but not *C5ar1* (CD88) (**Figure SVB**), suggesting that these cells likely represent bona fide cDC2. Recently, *Fscn1^+^Ccr7^+^* “mature DCs enriched in immunoregulatory molecules” (mReg-DC) have been described in mouse and human lung cancer (36). We analyzed whether *Fscn1^+^Ccr7^+^* mature DCs from atherosclerotic aortas share features with this mReg-DC transcriptomic signature. Compared to cDC1 and cDC2 populations, aortic mature *Fscn1*^+^*Ccr7*^+^ DCs were clearly enriched for several genes characteristic of the mReg-DC signature, such as *Il4i1, Cd274, Tnfrsf4*, or *Ccl22*, and transcripts encoding co-stimulatory molecules (*Cd40, Cd80, Cd86*) (**Figure SVA**).

### Integrated analysis of human macrophages in atherosclerosis uncovers three major macrophage populations

To gain further insight into the transcriptional state of macrophages in human atherosclerosis, we integrated scRNA-seq data from Fernandez et al. investigating carotid endarterectomy specimens (24), and Wirka et al. analyzing atherosclerotic coronary arteries from explanted hearts of transplant recipients (17). Data from Fernandez et al.(24) were obtained from batch corrected gene expression matrices, as provided by the authors, of 1 lesion analyzed by CITE-seq (Fernandez_CITE, n=254 cells), and plaques from 6 patients analyzed by scRNA-seq (4 asymptomatic patients: ASYM1 to 4, and 2 symptomatic patients: SYM1 and 2; n= 746 total cells), (**Figure SI**). Wirka et al. reported significant batch effects across patients (17), so that cells from each patient were considered as independent samples (referred to as Wirka_1 to Wirka_4) in computational integration analyses (**Figure SI**). Data from Wirka et al.(17) contained not only immune cells but all vascular cell types, so that we first identified mononuclear phagocytes (expressing e.g. *CD14, CD68, CSF1R, C1QA, CD52*), and extracted the corresponding data for further integration.

A total of 2,890 cells were included in the integrated analysis, and 10 clusters were recovered (**Figure 3A**). Macrophages were identified based on the expression of markers such as *CD68, C1QA* and *C5AR1* (**Figure 3B**). Differential gene expression analyses across clusters identified 3 major human (h) macrophage populations: hInflammatory-Mφ (*CD74*, *HLA-DRB1*), putative hFoamy-Mφ (*APOC1, APOE, FABP5, FABP4*), and hLYVE1-Mφ (*LYVE1, LGMN, MARCO*) (**Figure 3A and C**). Additional minor populations were characterized by expression of e.g. *C3*, *JUN* and *CCL4* (hC3-Mφ) and Type I IFN response macrophages (hIFNIC-Mφ cluster; *ISG15, IFI6, MX1*). In addition, cells corresponding to monocytes (hMonocytes: *VCAN, CD52, S100A8, S100A9, LYZ)(37*), were readily observed. Other minor clusters of proliferating cells (hProlif cluster: *TUBB*, *H2AFZ, STMN1*) and B cells (hB_cell cluster: *MZB1, JCHAIN*) were also observed (**Figure 3A and C**).

**Figure 3:**
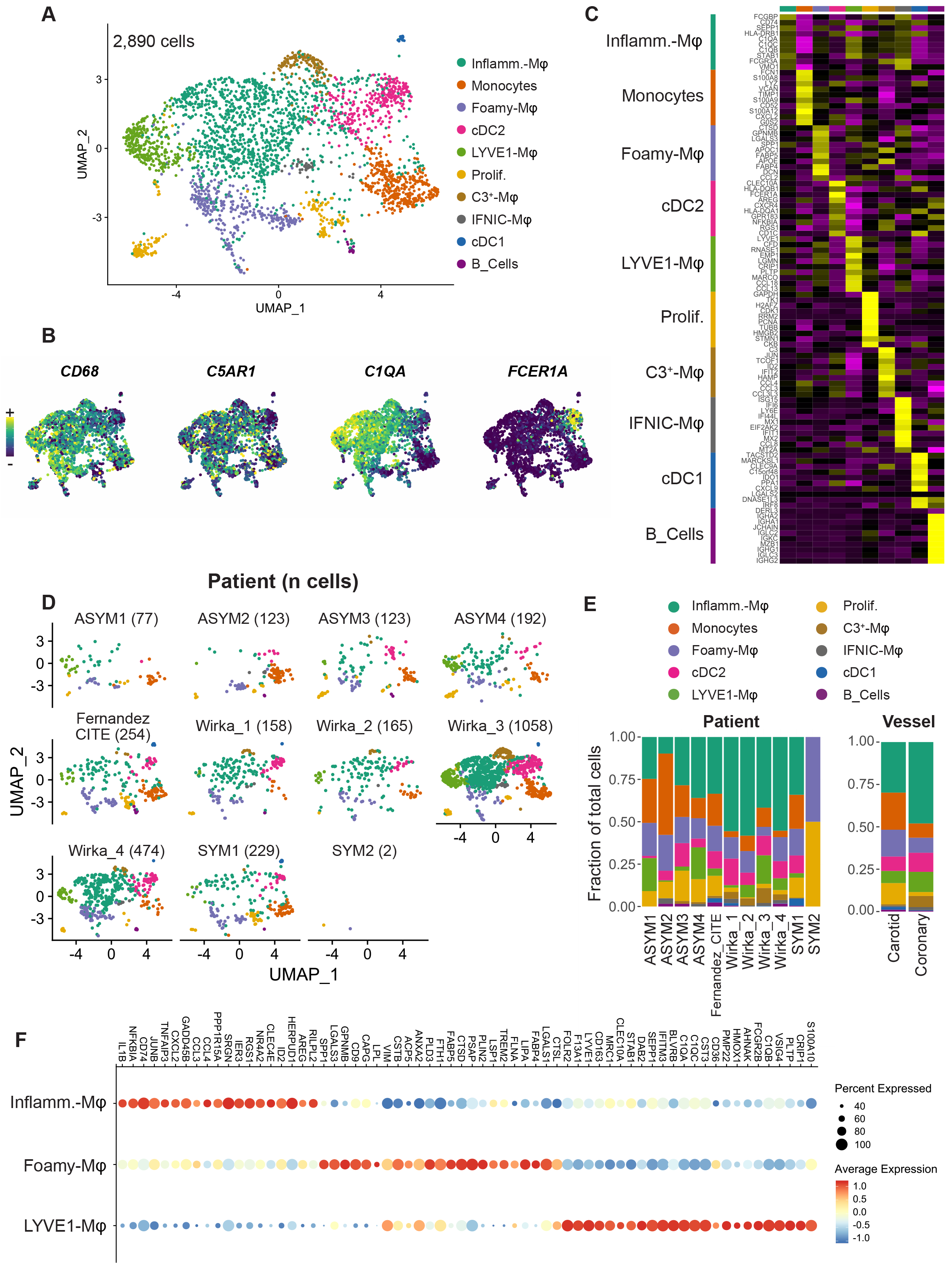
integrated scRNA-seq analysis of macrophages in human atherosclerosis. **A)** UMAP representation and clustering analysis of integrated scRNA-seq gene expression data in 2,890 human mononuclear phagocytes from atherosclerotic lesions; **B)** expression of C1QA and CD68 projected onto the UMAP plot; **C)** heatmap of averaged gene expression (top 10 genes ordered by fold change) in the clusters (Inflamm.=Inflammatory); **D)** projection of single cells in the UMAP space according to patient of origin; **E)** fraction represented by each cluster within total cells from each patient (left) or vascular bed (right); **F)** DotPlot showing the expression of transcripts enriched in human Inflammatory-Mφ, Foamy-Mφ and LYVE1-Mφ that are also enriched in their putative mouse counterparts (i.e. mouse Inflammatory-Mφ, Foamy/*Trem2*^hi^-Mφ, and Resident/Resident-like-Mφ respectively).

We could also recover cDC1 (hcDC1: *CLEC9A, IRF8, IDO1*) and cDC2 (hcDC2: *CLEC10A, FCER1A, CD1C*) (38) populations (**Figure 3A-B**). No cluster of mature DCs was readily observable, but we recovered a population of DCs with some features of the mature/mReg-DC signature (*CCL17, CCL19, MARCKSL1*, *IDO1*) when manually gating FSCN1^+^CCR7^+^ cells (n=9 cells) within total DCs (**Figure SVC and D**). Although these results need to be cautiously interpreted given the low number of cells analyzed, they suggest that dendritic cell populations proximal to those observed in mice populate human atherosclerosis.

The number of analyzed cells per patient ranged from n=2 (patient SYM2 from ref.(24)) to 1053 (patient 3 from (17)) (**Figure 3D**). Proportions of cell clusters varied greatly across patients (**Figure 3E**), which may reflect the well-known heterogeneity of plaque composition and morphology in patients (39–41) and difficulty in retrieving cells from human atherosclerotic tissues, and stresses the need for additional studies including more patients and cells. Low number of cells in some patients (ASYM1, SYM2) precludes interpretation of cluster repartition in these samples (**Figure 3E**). All the clusters were present in both vascular beds (**Figure 3E**).

Gene expression patterns within the three main human macrophage populations (hInflammatory-Mφ, hFoamy-Mφ and hLYVE1-Mφ) clearly suggested proximity to the major mouse aortic macrophage subsets we previously identified (6). Thus, we next performed differential gene expression analysis specifically within the 3 main human macrophage clusters, namely hInflammatory-Mφ, hFoamy-Mφ and hLYVE1-Mφ, and examined overlap with marker genes of mouse aortic Inflammatory, foamy/*Trem2*^hi^ and resident/resident-like macrophages. This revealed that besides MHCII encoding genes (*CD74*), hInflammatory-Mφ were enriched in inflammatory cytokines (*CXCL2, CCL3, CCL4, IL1B*), receptors (*CLEC4E)(42*) and transcriptional regulators (*IER3, NFKBIA, NR4A2*) similarly found in mouse Inflammatory-Mφ. hFoamy-Mφ showed an enrichment in markers characteristic of mouse foamy/*Trem2*^hi^ macrophages (*TREM2, CD9, GPNMB, SPP1, CTSL, LIPA, ACP5*), and hLYVE1-Mφ expressed genes associated with mouse resident/resident-like macrophages (*LYVE1, CD163, SEPP1, FOLR2, F13A1, MRC1, VSIG4*) (**Figure 3F**).

To further evaluate the similarity between macrophage clusters observed in human and mouse atherosclerosis, we performed gene ontology enrichment (GO) analyses, which revealed enrichment in similar biological processes, cellular components or molecular functions (**Supplementary Excel file**). In particular, hFoamy-Mφ and mouse foamy/*Trem2*^hi^ macrophages were enriched for putative functions related to lipid metabolism (e.g. biological process GO terms: lipid catabolic process, lipid storage, cellular response to lipoprotein particle stimulus), and similar molecular functions (e.g. molecular function GO terms: fatty acid binding, lipase activity, antioxidant activity, low-density lipoprotein particle binding) (**Supplementary Excel file**).

### Cross species integration reveals conserved macrophage transcriptional states in mouse and human atherosclerosis

To further investigate the proximity of the transcriptional states of mouse to human macrophages in atherosclerosis, we performed cross-species integration of scRNA-seq data. Mouse gene symbols were converted to their human homologs using the BioMart-Ensembl database. Mouse datasets were pre-processed to identify and extract cells corresponding to mononuclear phagocytes (macrophages, monocytes, dendritic cells), and integrated with the human data. After integration in Seurat v3 and dimensional reduction, clustering analysis generated 10 clusters (**Figure 4A**), with a clear mouse to human overlap (**Figure 4B**). By identifying and annotating cell clusters based on the characteristic gene expression patterns identified in the integrated human data and based on the mouse homologues in the integrated mouse data, we could readily recover integrated (int) Inflammatory macrophages (int-Inflammatory-Mφ: *CD83, CCRL2, IFRD1*), int-Res/Res-like-Mφ (*LYVE1, FOLR2, F13A1*), int-Foamy/TREM2hi-Mφ (*GPNMB, CD9, SPP1, FABP5*), int-MoDC/cDC2 (*NAPSA*, *KLRD1*), int-Monocytes (*PLAC8, MSRB1, THBS1*), int-IFNIC-Mφ (*IRF7*, *ISG15*), int-*FSCN1*/*CCR7*-DCs, and int-*XCR1/IRF8*-cDC1 (**Figure 4C**). All clusters contained both mouse and human cells (**Figure 4B**). This suggests that cDC1, cDC2, mature DCs, classical monocytes, and macrophages observed in mouse atherosclerotic aortas are also found within human lesions, and that macrophages that populate human lesions display transcriptional states proximal to the major mouse aortic macrophage subsets (Resident/Resident-like, Inflammatory, Foamy/Trem2^hi^, IFNIC).

**Figure 4:**
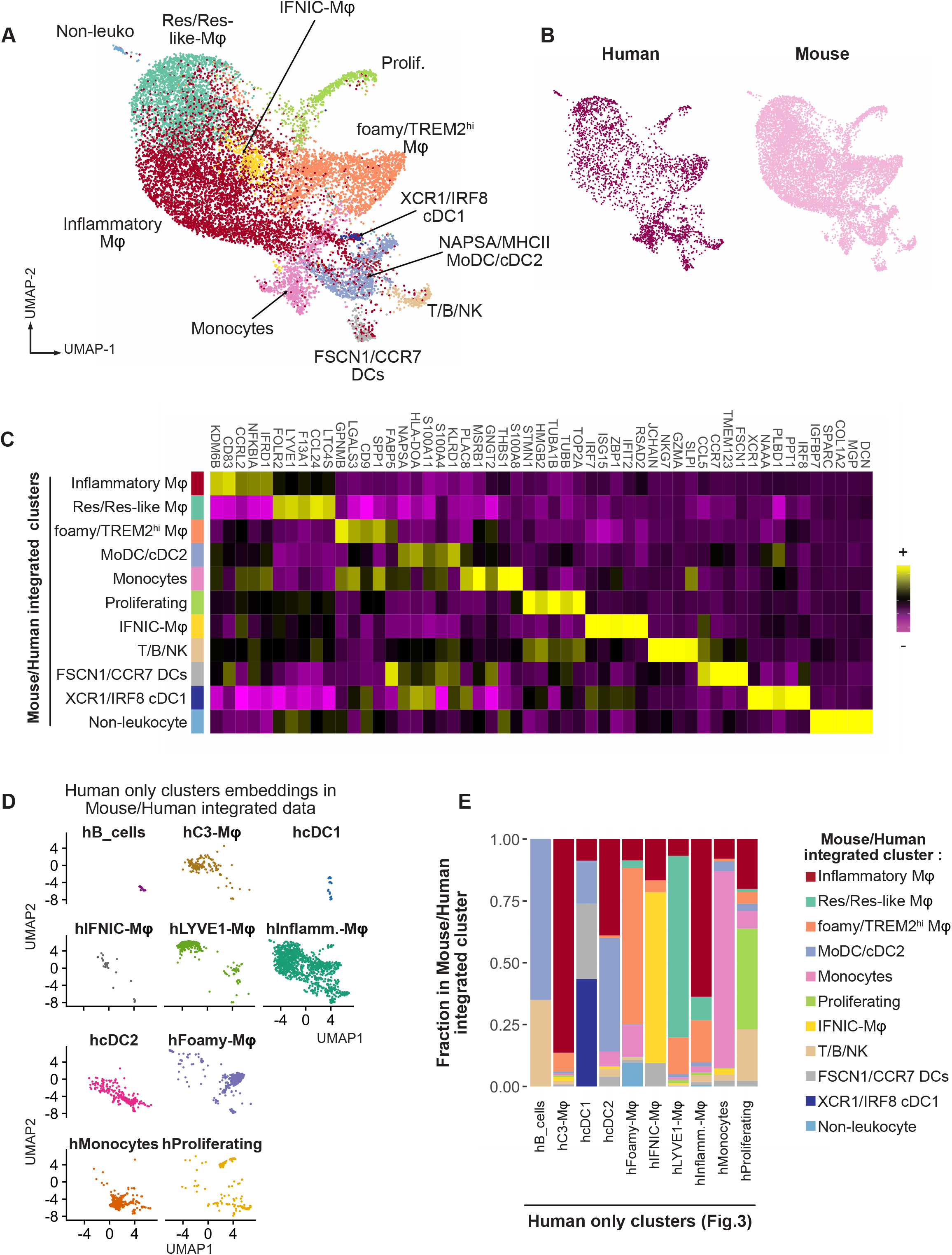
Cross-species scRNA-seq integrated analysis of macrophages in murine and human atherosclerosis. **A)** UMAP plot and clustering analysis (with annotation) of the mouse/human integrated data; **B)** cell species of origin projected onto the UMAP plot; **C)** heatmap of enriched genes in the integrated cluster (top 5 ordered by fold change); **D)** projecton of the human-only clusters (see Figure 2) on the UMAP plot of the mouse/human integrated data; **E)** mapping of the human-only clusters in the mouse/human integrated clusters.

We further mapped the UMAP-embeddings and clustering characteristics of the cellular states defined in the human data (see **Figure 3A**) to the mouse/human integrated data (**Figure 4A**). Overall, the major human macrophage subtypes mapped to the expected integrated clusters (**Figure 4D and E**). hInflammatory-Mφ mapped mostly to int-Inflammatory-Mφ (64%), hFoamy-Mφ mapped to int-Foamy/TREM2hi-Mφ (63%), hIFNIC-Mφ to int-IFNIC-Mφ (69%), hLYVE1-Mφ to int-Res/Res-like-Mφ (73%) hMonocytes to int-Monocytes (80%), and hC3^+^-Mφ to int-Inflammatory-Mφ (86%) (**Figure 4E**). However, we did not observe a full overlap between human and integrated clusters (e.g. 17% of hInflammatory-Mφ mapped to int-Foamy/TREM2hi-Mφ), which could be attributed to macrophages with intermediate transcriptional profiles between two polarized states, and loss of information due to inclusion of only those genes with unambiguous human-to-mouse homologs in the cross-species integrated analysis. This was also particularly clear for other cell populations, such as human cDC2, which mapped almost equally to int-MoDC/cDC2 (46%) and int-Inflammatory-Mφ (39%). This can be explained by shared enrichment for inflammatory markers (e.g. *IL1B*, *SOCS3*), and specific markers of human cDC2 not being conserved (*CD1C*) or showing a wider/distinct expression patterns (*FCER1A, CLEC10A*) in mice. Altogether, this cross-species integration analysis substantiates the notion that macrophages with similar transcriptional states populate human and mouse atherosclerotic lesions.

## Discussion

We here show that major macrophage and dendritic cell transcriptional states are conserved across widely employed mouse models of atherosclerosis, and that human lesions are populated by mononuclear phagocytes displaying transcriptional states resembling those found in mouse atherosclerosis. We provide two layers of evidence for this conclusion: (i) integrated analysis of human lesion scRNA-seq data from two independent studies recovered macrophage and dendritic cell states highly proximal to those observed in mice and (ii) direct cross-species data integration showed a strong overlap between mouse and human mononuclear phagocyte states.

Accumulation of macrophage foam cells in the intima is instrumental to lesion development. Macrophages reminiscent of the foamy/*Trem2^hi^* macrophage state were observed across mouse models of atherosclerosis and in human lesions. Mononuclear phagocytes with a transcriptional signature proximal to the foamy/Trem2^hi^ macrophage state found in atherosclerosis have been observed in mouse models of neurodegenerative disease (disease associated microglia, DAM (43)), demyelinating disease (44), non-alcoholic steatohepatitis (NAM: NASH associated macrophages) (45), metabolic-associated fatty liver disease (46), liver fibrosis (SAM: scar associated macrophages) (47), and diet-induced obesity (LAM: lipid-associated macrophages) (48). In the diseased liver and adipose tissue, features of this transcriptomic state were similar in mice and humans, with many transcriptomic or cell surface markers such as *TREM2, SPP1* or *CD9* (45, 47), (46) being conserved across species. The situation seems more complex in neurodegeneration-associated microglia as the characteristic DAM signature was not readily detected in single-nucleus RNA-seq (snRNA-seq) analysis of human neurodegenerative brain samples (49). However, this observation might be due to technical issues, as characteristic genes of the DAM signature such as *APOE* or *SPP1* were poorly detected in single-nucleus compared to single-cell RNA-seq (50). This technical limitation, if further confirmed, needs to be taken into consideration in future snRNA-seq analyses of atherosclerotic samples.

LAM, DAM and atherosclerosis-associated foamy/*Trem2*^hi^ macrophages share expression of a set of genes with enrichment in lipid metabolism pathways (48), suggesting that similar mechanisms related to lipid loading may drive acquisition of this macrophage state. Nevertheless, further analyses will be required to determine the fine tissue- and species-specific particularities of these macrophages. Evidence from neurodegenerative disease models indicate that acquisition of the DAM state may depend on *Apoe* (51), which raised the possibility that acquisition of the foamy/Trem2^hi^ macrophage state might differ between the most widely employed mouse models of atherosclerosis, i.e *Ldlr^-/-^* and *Apoe^-/-^* mice. However, consistent with previous reports (6, 7), our integrated analysis indicates that *Apoe* expression appears dispensable for the acquisition of the foamy/*Trem2*^hi^ macrophage state in mouse arteries. Fully elucidating the impact of the mouse genotype on macrophage states will require more suitable experimental designs including biological replicates and differential gene expression in the absence of overt batch correction, e.g. by employing single-cell multiplexing technologies such as cell hashing (52) or MULTI-Seq (53). Recently, we identified *Trem2^hi^* macrophages in the ischemic mouse heart sharing gene expression similarities with the LAM/DAM/foamy signature (28), and two reports identified *Trem2* enriched immunosuppressive macrophages in tumor models (54), (55), indicating that part of this transcriptional signature may not only be related to pathological lipid loading, but rather more generally induced in contexts of tissue damage. Our observation that macrophages with a *Trem2^hi^* signature populate the aorta in the context of Angiotensin-II mediated inflammation corroborates this notion. Major transcriptional hubs involved in regulation of lipid homeostasis such as the liver-X-receptor (LXR) pathway are activated also in response to efferocytosis of dead cells (56), raising the possibility that macrophages with high efferocytic activity may also acquire a foamy/*Trem2^hi^* gene expression signature.

By analyzing a large number of mouse macrophages in our integrated approach, we could identify discrete subpopulations within foamy/*Trem2^hi^* macrophages, including recently identified aortic intimal resident macrophages (MAC-AIR) (15). We furthermore identified two subsets we termed *Trem2^hi^-Slamf9* and *Trem2^hi^-Gpnmb*. Analysis of gene expression patterns is in line with the notion that MAC-AIR present a gene expression signature specific to the vascular intimal niche that is acquired by infiltrating monocytes, and that lesion-associated foamy macrophages further attain the expression of a disease specific gene signature (15). Importantly, canonical markers of this signature (*Trem2, Spp1*) appeared to have a low expression in MAC-AIR. While strong recovery of these three clusters in single-cell data from aortic intimal foamy cells (7) and recent fate-mapping and imaging analysis (15) clearly suggest that these populations populate the intima, their precise localization within the complex morphology of arteries remains to be further defined. Likewise, while Williams et al. have identified MAC-AIR as self-renewing resident macrophages seeded by monocytes during the perinatal period (15), the ontogeny of *Trem2^+^-Slamf9* and *Trem2^hi^-Gpnmb* macrophages requires further investigation. However, extrapolation of decades of research on foamy macrophage accumulation in atherosclerosis clearly suggests a monocytic origin of these cells, which could be confirmed in future analyses using recently developed monocyte fate-mapping models based on *Ms4a3* (57) or *Cxcr4* (58). Finally, whether similar subpopulations of *foamy/Trem2^hi^* macrophages can be detected in human lesions will require sampling of a larger number of human lesional macrophages.

We observed macrophages corresponding to inflammatory macrophages both in mice and humans. Compared to other macrophages, these are characterized by expression of genes encoding MHCII/HLA genes, and inflammatory cytokines (e.g. *IL1B*, inflammatory chemokines). Macrophages with a type I interferon signature (IFNIC)(14) were also observed in mice and humans.

In mice, resident/resident like macrophages are defined by expression of a characteristic set of genes (*Lyve1, Folr2, Sepp1, F13a1, Pf4, Cd163*). In human lesional macrophages, we identified a subset of cells with a clearly overlapping transcriptional state. The exact localization of these cells in human diseased vessels remains to be fully elucidated. In mice, Lyve1 ^+^ macrophages are typically located in the adventitia (12, 13). We had previously observed LYVE1 protein expression in macrophages in carotid endarterectomy specimens by immunohistochemistry (6), and we here detected *LYVE1^+^* macrophages in scRNA-seq in carotid endarterectomy plaques. As the adventitia is not extracted during the carotid endarterectomy procedure, this indicates that cells with the Resident/Resident-like/LYVE1^+^ state may be found within human lesions. However, in a recent report, Alsaigh et al. performed single-cell analysis of atherosclerotic lesions from 3 patients, where cells from the atherosclerotic core and the proximal adjacent coronary artery were analyzed (59). Macrophages with the foam cell signature (*APOE*, *APOC*1) were observed in the atherosclerotic core, while LYVE1 enriched macrophages were in the proximal adjacent coronary artery (59). Future investigations employing spatial transcriptomic methods (60) will help shed light on the precise localization of macrophage populations within diseased vessels.

While our work provides proof-of-concept of conserved transcriptional features of macrophage states across mouse and human atherosclerosis, further acquisition of high-quality data is clearly needed to fully elucidate the phenotypical landscape of human lesional macrophages, and cross-species characteristics of macrophage states. Altogether, our analysis encompasses cells from 11 patients across 2 vascular beds (carotid and coronary arteries). While mice employed in experimental models of atherosclerosis are rather homogeneous (same genetic background, age, sex, absence of additional comorbidities), patients represent a highly heterogeneous population, and many factors (e.g. age, sex, comorbidities such as diabetes, etc…) are known to influence plaque immune composition (20, 61–63). Even in patients with a similar clinical profile, atherosclerotic lesions are highly heterogeneous in their morphology and cellular composition (39–41). Hence, we propose that to identify macrophage transcriptional states correlated to clinically relevant events, analysis of a vast number of cells from many patients, and of cells from different vascular beds, will be necessary. Besides increasing statistical power to balance patient and plaque heterogeneity, technical issues remain an additional important hurdle for single-cell analyses of clinical samples (64). The human and mouse studies included here all employed enzymatic digestion of tissues, which leads to cell recovery that may not represent the true composition of *in vivo* lesions as some cells (in particular macrophage foam cells) may be more difficult to recover compared to other cells, and which causes artificial gene expression patterns such as induction of immediate early genes (27, 65). Single-nucleus RNA-seq, which bypasses the need for enzymatic tissue digestion, might be particularly suited to the analysis of human atherosclerosis, although poor detection of key mononuclear phagocyte genes may need to be carefully accounted for (50).

Altogether, our work provides proof of concept that macrophage transcriptomic states in atherosclerosis are conserved across mouse models of the disease and different vascular beds in humans. These findings are of importance for experimental investigations of macrophage function in atherosclerosis and their potential clinical translation. However, further research is critically needed to obtain a better understanding of macrophage populations and their states in human atherosclerosis, and of the fine differences between the human and mouse species that may bias pre-clinical investigations. Such research will benefit both from methodological advances as well as the analyses of substantially increased numbers of patients.

## Acknowledgements

A.Z. was supported by the Interdisciplinary Center for Clinical Research (Interdisziplinäres Zentrum für Klinische Forschung [IZKF]), University Hospital Würzburg (E-352 and A-384), and the Deutsche Forschungsgemeinschaft (DFG; German Research Foundation, 374031971 - TRR 240, 324392634 - TR221, ZE827/13-1, 14-1, 15-1, and 16-1). This study was further supported by the SFB 1123 project A07 (to C.S.), as well as the DZHK (German Centre for Cardiovascular Research) and the BMBF (German Ministry of Education and Research) (grants 81Z0600204 to C.S. and 81X2600252 to T.W.), and the BMBF within the Comprehensive Heart Failure Centre Würzburg (BMBF 01EO1504 to C.C. and A-E.S.). K.L. was supported by NIH P01 HL136275, R35 HL145241, and R01 HL146134. A.-E.S. is supported by the EMBO Young Investigator Program. C. C. was supported by the Interdisciplinary Center for Clinical Research (IZKF), University Hospital Würzburg (E-353) and the DFG (CO1220/1-1).

---

**Table S1.**
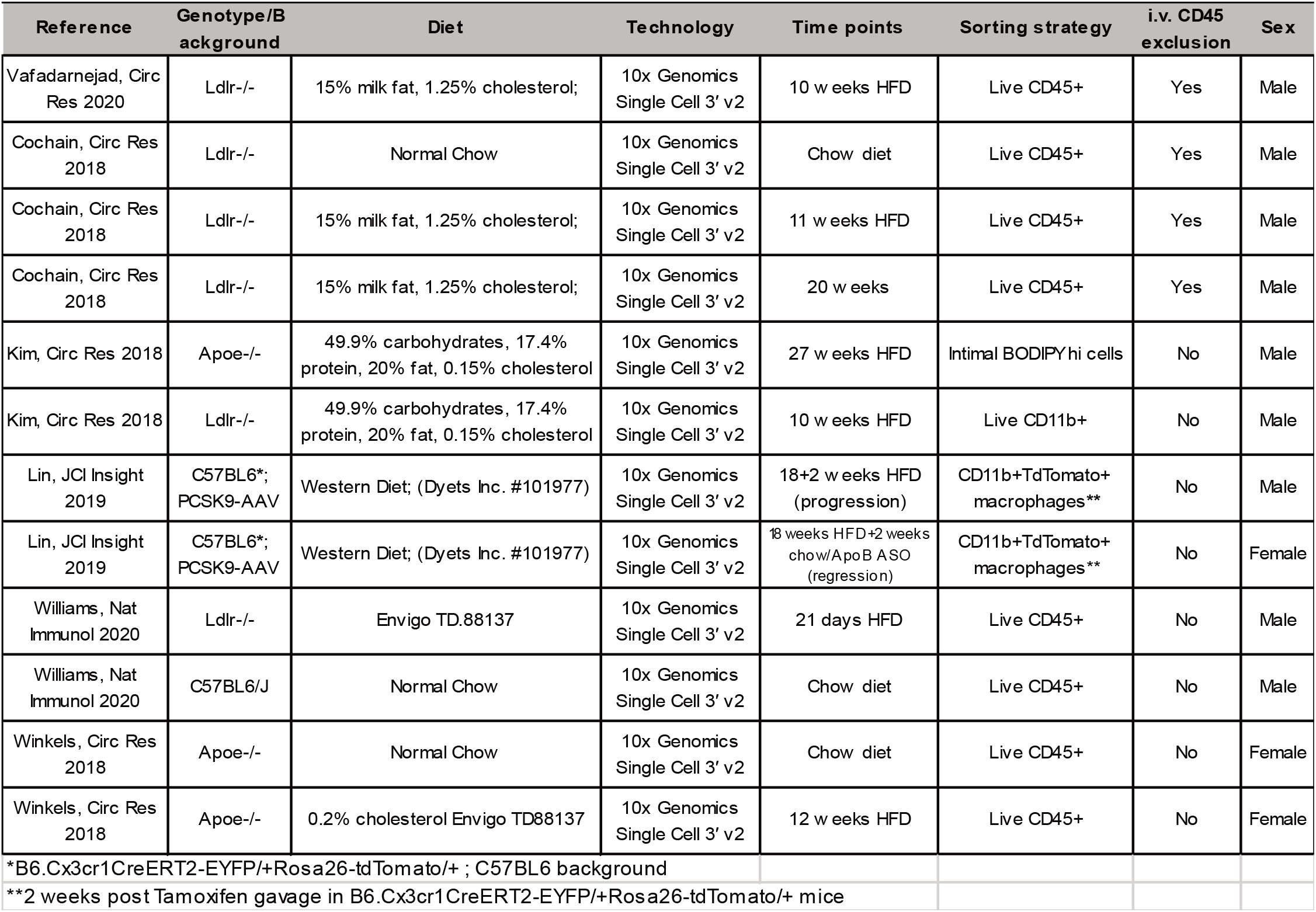

**Figure SI:**
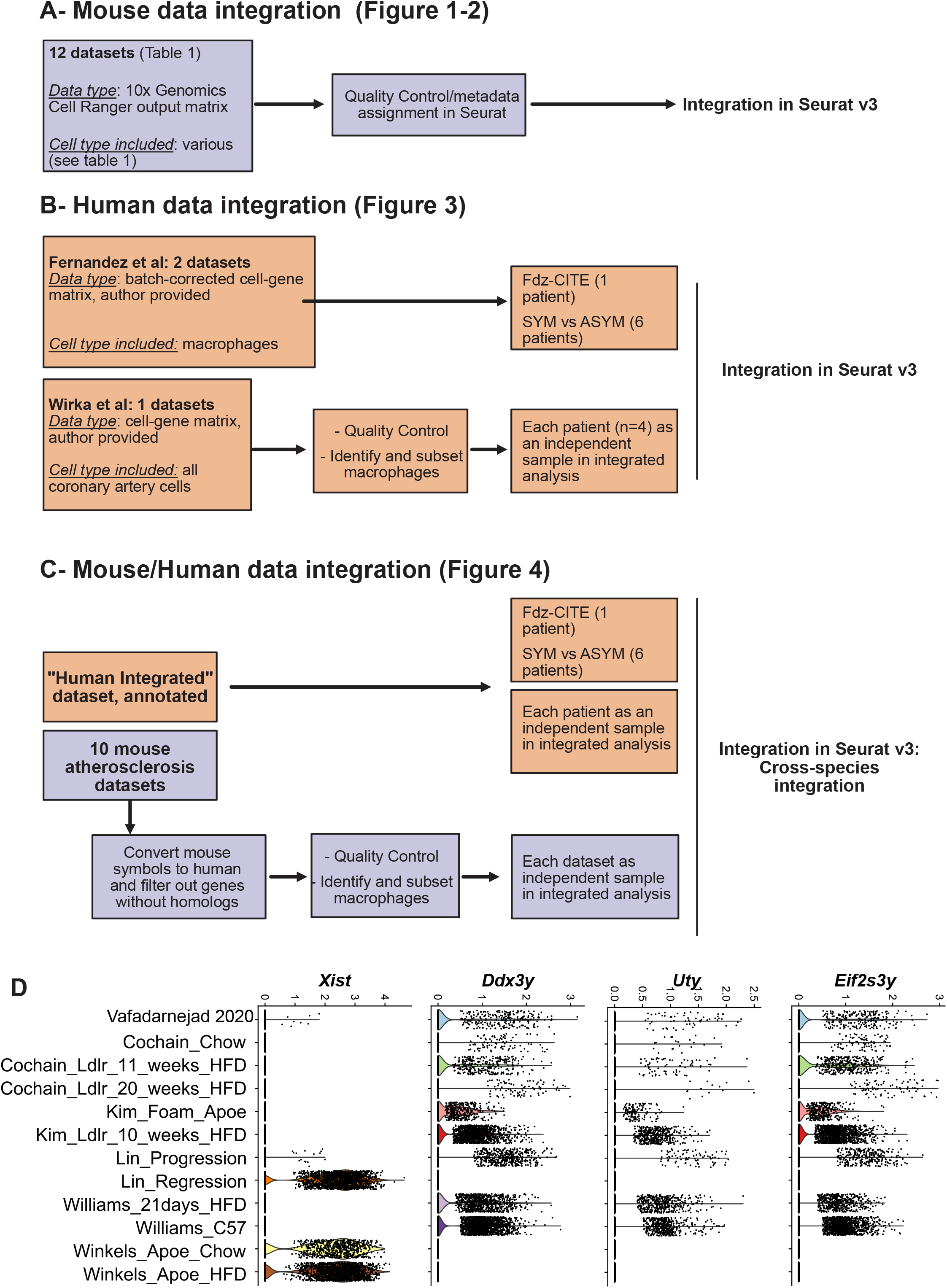
A-C schematic representation of the integrated analysis procedure and D) analysis of sex specific gene expression in mouse single-cell studies.

**Figure SII:**
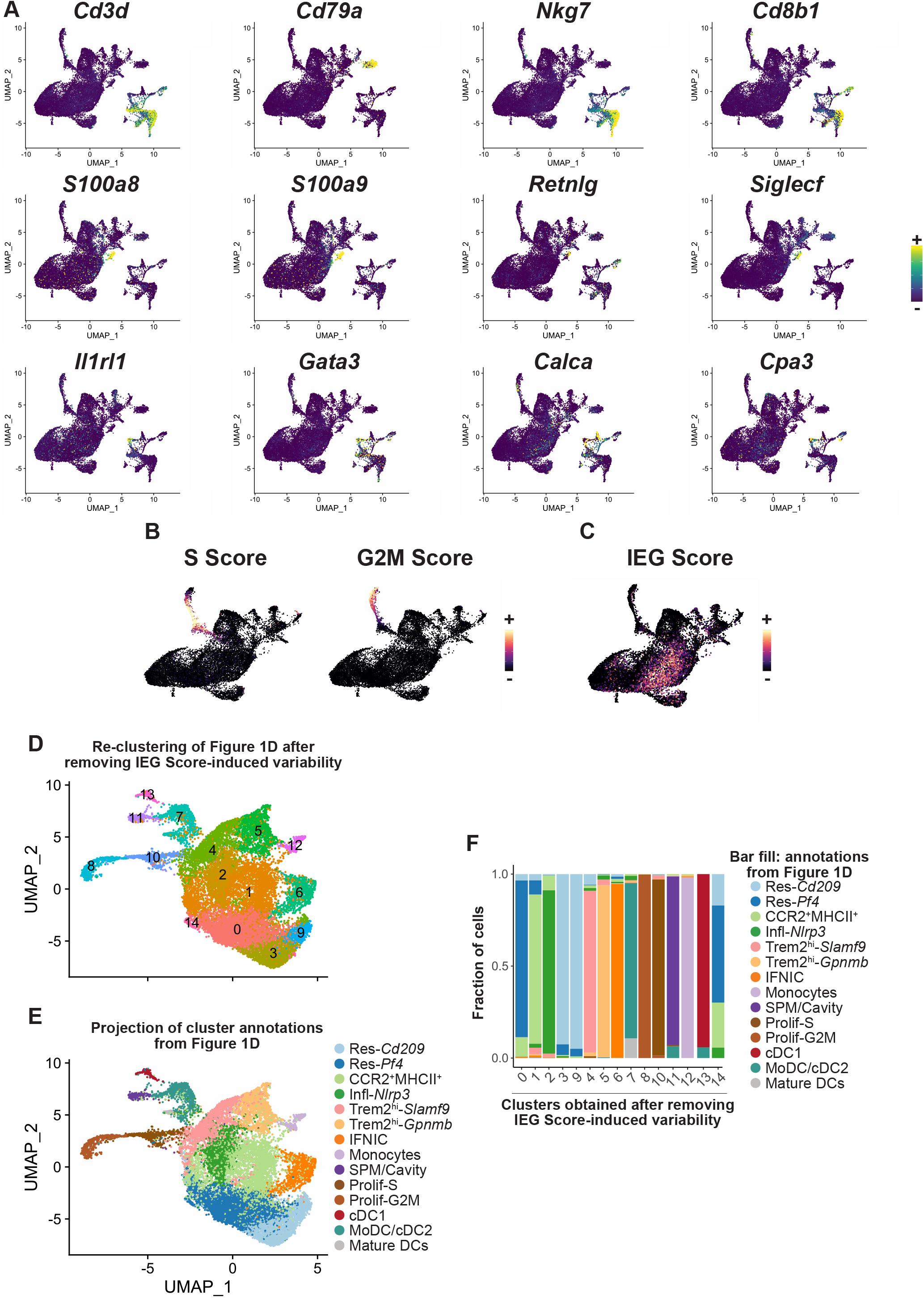
Identification of immune cell lineages, cell cycle phase and immediate early genes expression in aortic immune cells (related to Figure 1). **A)** expression of the indicated transcripts identifying immune cell lineages projected onto the UMAP plot shown in Figure 1A; **B)** S and G2M cell cycle score and **C)** Immediate Early Gene (IEG) expression score in mononuclear phagocytes projected onto the UMAP plot. **D-F)** re-clustering of mononuclear phagocytes after regressing out gene expression variability caused by IEG expression with **D)** UMAP plot of the new clustering analysis, **E)** projection of clusters obtained in **Figure 1D** on the new UMAP embeddings and **F)** distribution of the clusters obtained in Figure 1D across the clusters obtained after removing IEG Score-induced variability.

**Figure SIII:**
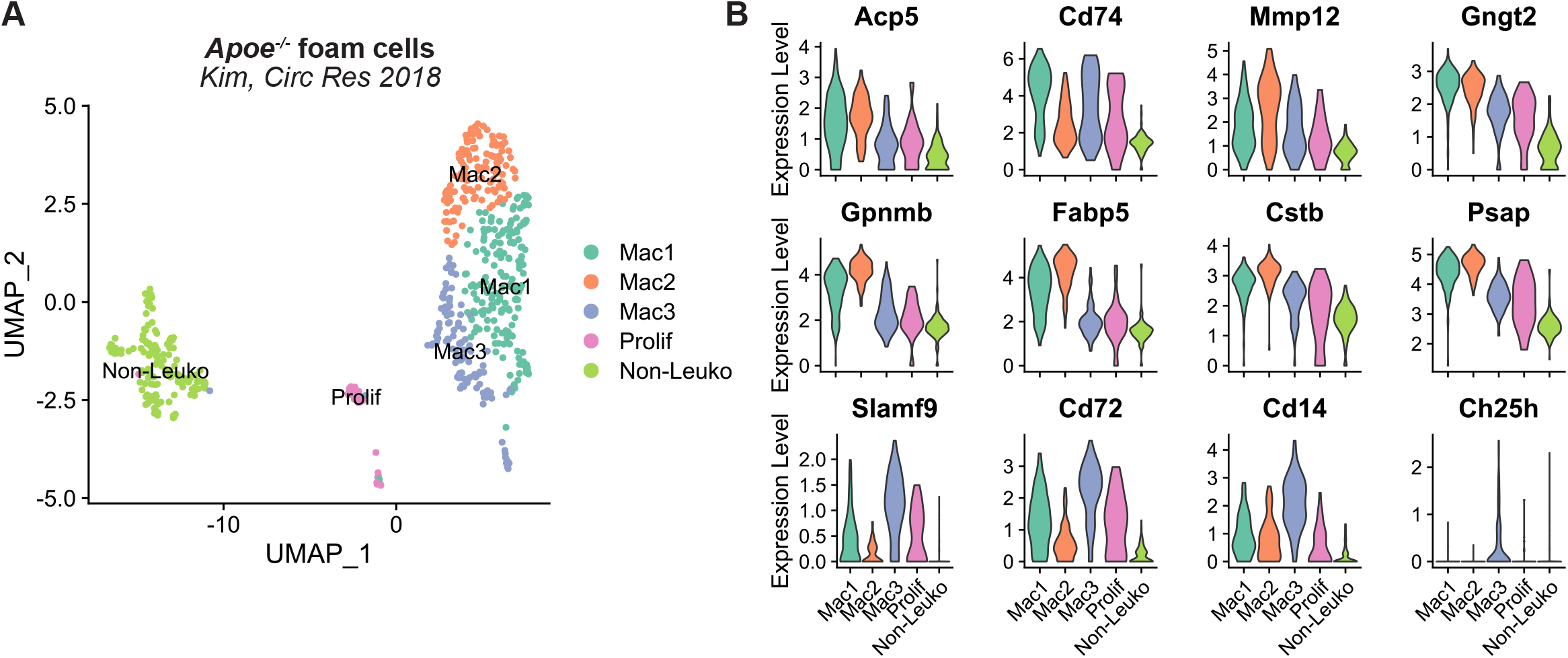
Analysis of scRNA-seq data from the *Apoe*^-/-^ Foam Cell (Kim, 2018) data set alone (related to Figure 2). **A)** UMAP plot and cluster annotation (Mac=macrophage; Prolif: proliferating; non-leuko=non leukocyte) and **B)** expression of the indicated transcripts in the different clusters (prolif=proliferating)

**Figure SIV:**
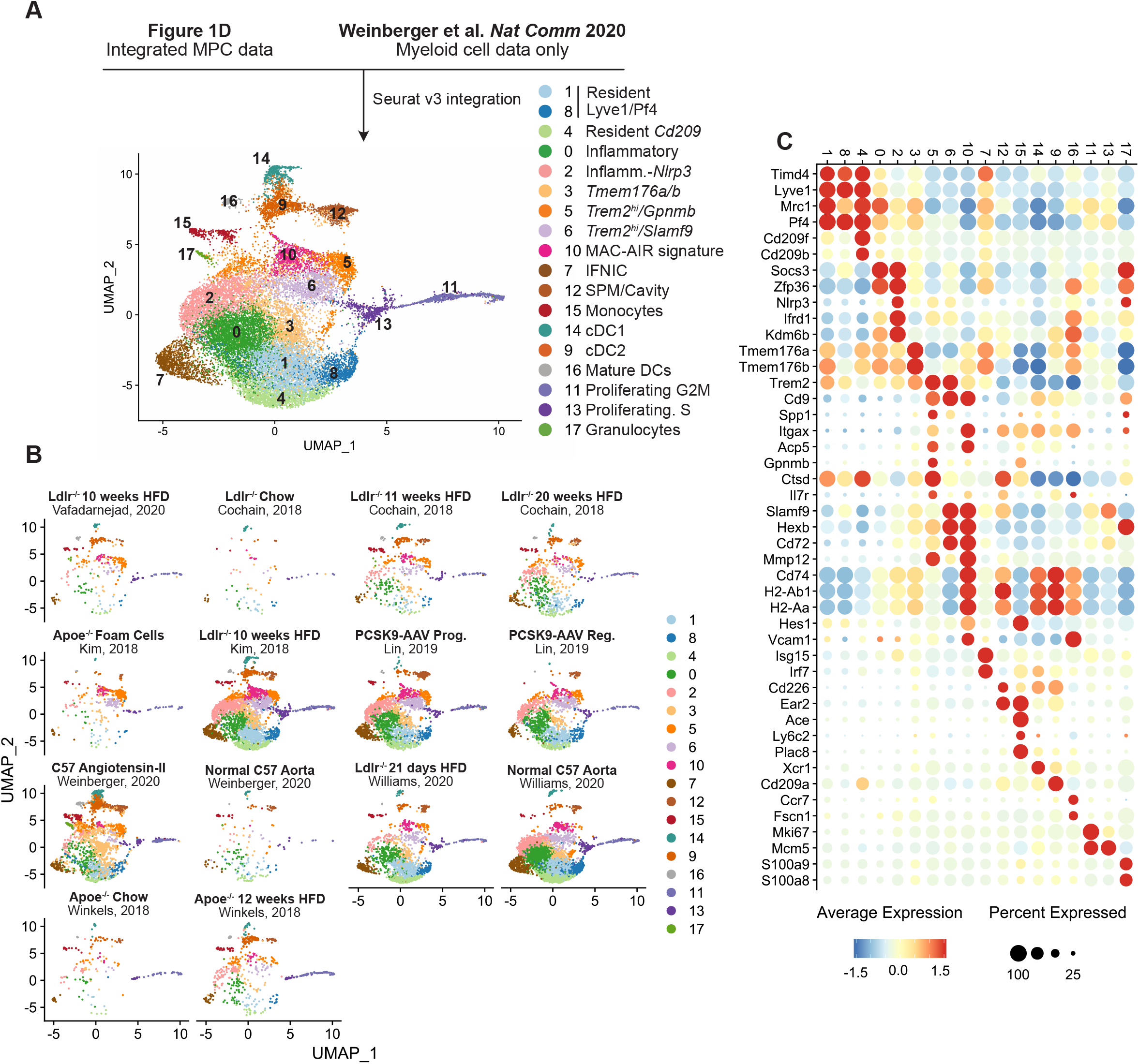
Identification of mononuclear phagocyte subsets in AngII induced aortic inflammation (related to Figure 1 and 2). The integrated mononuclear phagocyte dataset (Figure 1D) was integrated with myeloid cell scRNA-seq data from Weinberger et al. Nature Communications 2020. **A)** UMAP plot with color-coded identification of clusters and **B)** UMAP-plot splitted by dataset of origin; **C**) dot plot showing average expression of selected marker genes in the clusters.

**Figure SV:**
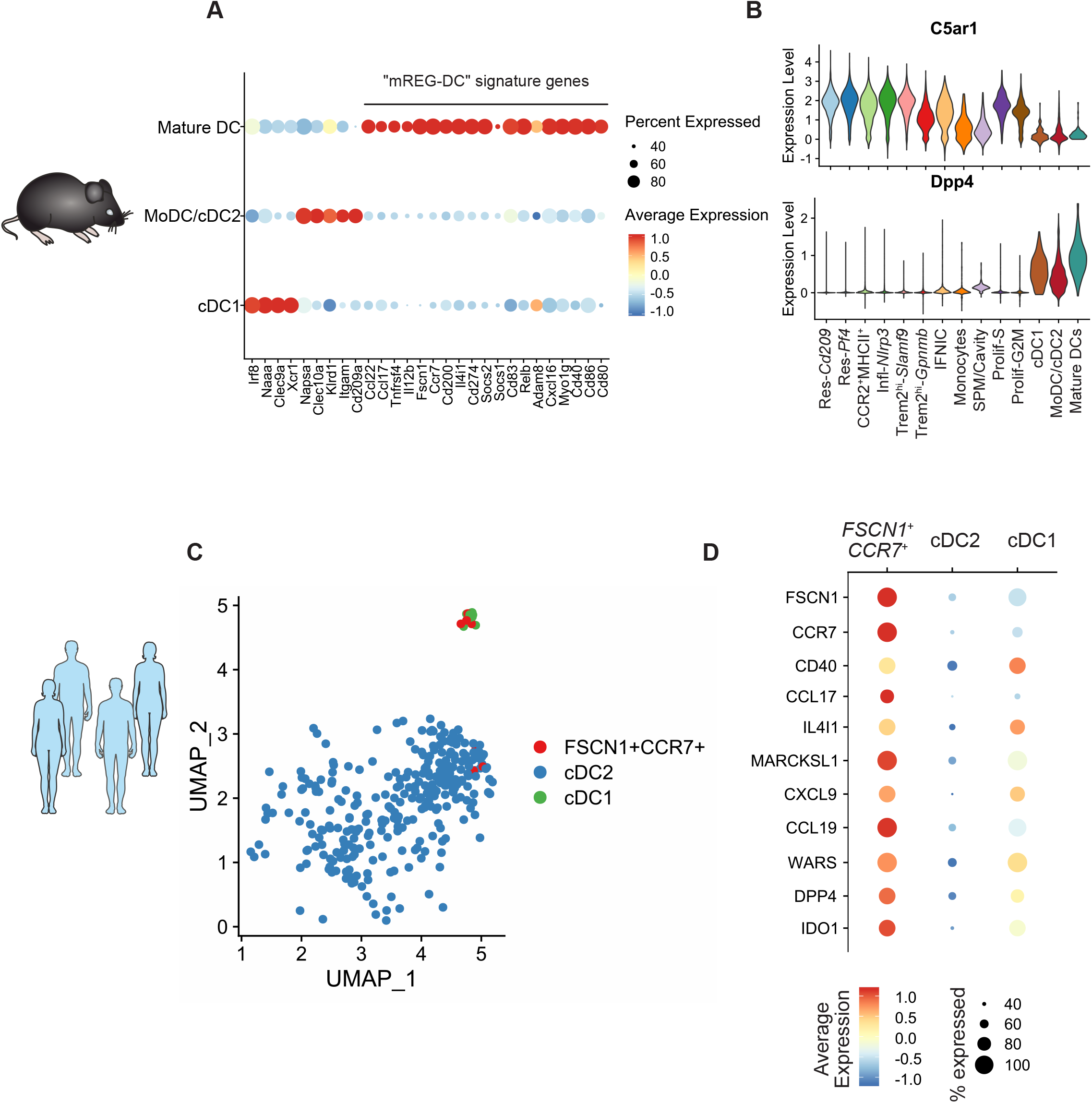
Identification of the mREG-DC signature in mouse and human atherosclerosis. **A)** Gene expression DotPlot in the 3 dendritic cell populations shows expression of “mregDC” signature genes in mature DCs.**B)** Violin plots showing expression of *Dpp4* (encoding CD26) and *C5ar1* (encoding CD88) in the mononuclear phagocyte subsets. **C)** Cells coexpressing *FSCN1* and *CCR7 (FSCN1^+^CCR7^+^* DCs, n=10 cells) were manually selected in Seurat and are color coded on the UMAP plot (same coordinates as Figure 3A, only DC populations shown). **D)** Dot Plot showing average expression and proportion of expressing cells for the indicated transcripts in *FSCN1^+^CCR7^+^* DC, cDC2 and cDC1 populations.

## References

1. Libby P, Buring JE, Badimon L, Hansson GK, Deanfield J, Bittencourt MS, Tokgozoglu L, Lewis EF. 2019. Atherosclerosis. Nat Rev Dis Primers 5: 56

2. Wolf D, Ley K. 2019. Immunity and Inflammation in Atherosclerosis. Circ Res 124: 315–27

3. Cochain C, Zernecke A. 2017. Macrophages in vascular inflammation and atherosclerosis. Pflugers Arch 469: 485–99

4. Willemsen L, de Winther MPJ. 2020. Macrophage subsets in atherosclerosis as defined by single-cell technologies. J Pathol

5. Zernecke A, Winkels H, Cochain C, Williams JW, Wolf D, Soehnlein O, Robbins CS, Monaco C, Park I, McNamara CA, Binder CJ, Cybulsky MI, Scipione CA, Hedrick CC, Galkina EV, Kyaw T, Ghosheh Y, Dinh HQ, Ley K. 2020. Meta-Analysis of Leukocyte Diversity in Atherosclerotic Mouse Aortas. Circ Res 127: 402–26

6. Cochain C, Vafadarnejad E, Arampatzi P, Pelisek J, Winkels H, Ley K, Wolf D, Saliba AE, Zernecke A. 2018. Single-Cell RNA-Seq Reveals the Transcriptional Landscape and Heterogeneity of Aortic Macrophages in Murine Atherosclerosis. Circ Res 122: 1661–74

7. Kim K, Shim D, Lee JS, Zaitsev K, Williams JW, Kim KW, Jang MY, Seok Jang H, Yun TJ, Lee SH, Yoon WK, Prat A, Seidah NG, Choi J, Lee SP, Yoon SH, Nam JW, Seong JK, Oh GT, Randolph GJ, Artyomov MN, Cheong C, Choi JH. 2018. Transcriptome Analysis Reveals Nonfoamy Rather Than Foamy Plaque Macrophages Are Proinflammatory in Atherosclerotic Murine Models. Circ Res 123: 1127–42

8. Winkels H, Ehinger E, Vassallo M, Buscher K, Dinh HQ, Kobiyama K, Hamers AAJ, Cochain C, Vafadarnejad E, Saliba AE, Zernecke A, Pramod AB, Ghosh AK, Anto Michel N, Hoppe N, Hilgendorf I, Zirlik A, Hedrick CC, Ley K, Wolf D. 2018. Atlas of the Immune Cell Repertoire in Mouse Atherosclerosis Defined by Single-Cell RNA- Sequencing and Mass Cytometry. Circ Res 122: 1675–88

9. Lin JD, Nishi H, Poles J, Niu X, McCauley C, Rahman K, Brown EJ, Yeung ST, Vozhilla N, Weinstock A, Ramsey SA, Fisher EA, Loke P. 2019. Single-cell analysis of fate-mapped macrophages reveals heterogeneity, including stem-like properties, during atherosclerosis progression and regression. JCI Insight 4

10. Cochain C, Saliba AE, Zernecke A. 2018. Letter by Cochain et al Regarding Article, “Transcriptome Analysis Reveals Nonfoamy Rather Than Foamy Plaque Macrophages Are Proinflammatory in Atherosclerotic Murine Models”. Circ Res 123: e48–e9

11. Kim K, Choi JH. 2018. Response by Kim and Choi to Letter Regarding Article, “Transcriptome Analysis Reveals Nonfoamy Rather Than Foamy Plaque Macrophages Are Proinflammatory in Atherosclerotic Murine Models”. Circ Res 123: e50

12. Lim HY, Lim SY, Tan CK, Thiam CH, Goh CC, Carbajo D, Chew SHS, See P, Chakarov S, Wang XN, Lim LH, Johnson LA, Lum J, Fong CY, Bongso A, Biswas A, Goh C, Evrard M, Yeo KP, Basu R, Wang JK, Tan Y, Jain R, Tikoo S, Choong C, Weninger W, Poidinger M, Stanley RE, Collin M, Tan NS, Ng LG, Jackson DG, Ginhoux F, Angeli V. 2018. Hyaluronan Receptor LYVE-1-Expressing Macrophages Maintain Arterial Tone through Hyaluronan-Mediated Regulation of Smooth Muscle Cell Collagen. Immunity 49: 326–41 e7

13. Ensan S, Li A, Besla R, Degousee N, Cosme J, Roufaiel M, Shikatani EA, El-Maklizi M, Williams JW, Robins L, Li C, Lewis B, Yun TJ, Lee JS, Wieghofer P, Khattar R, Farrokhi K, Byrne J, Ouzounian M, Zavitz CC, Levy GA, Bauer CM, Libby P, Husain M, Swirski FK, Cheong C, Prinz M, Hilgendorf I, Randolph GJ, Epelman S, Gramolini AO, Cybulsky MI, Rubin BB, Robbins CS. 2016. Self-renewing resident arterial macrophages arise from embryonic CX3CR1(+) precursors and circulating monocytes immediately after birth. Nat Immunol 17: 159–68

14. King KR, Aguirre AD, Ye YX, Sun Y, Roh JD, Ng RP, Jr., Kohler RH, Arlauckas SP, Iwamoto Y, Savol A, Sadreyev RI, Kelly M, Fitzgibbons TP, Fitzgerald KA, Mitchison T, Libby P, Nahrendorf M, Weissleder R. 2017. IRF3 and type I interferons fuel a fatal response to myocardial infarction. Nat Med 23: 1481–7

15. Williams JW, Zaitsev K, Kim KW, Ivanov S, Saunders BT, Schrank PR, Kim K, Elvington A, Kim SH, Tucker CG, Wohltmann M, Fife BT, Epelman S, Artyomov MN, Lavine KJ, Zinselmeyer BH, Choi JH, Randolph GJ. 2020. Limited proliferation capacity of aortic intima resident macrophages requires monocyte recruitment for atherosclerotic plaque progression. Nat Immunol 21: 1194–204

16. Fernandez DM, Rahman AH, Fernandez NF, Chudnovskiy A, Amir ED, Amadori L, Khan NS, Wong CK, Shamailova R, Hill CA, Wang Z, Remark R, Li JR, Pina C, Faries C, Awad AJ, Moss N, Bjorkegren JLM, Kim-Schulze S, Gnjatic S, Ma’ayan A, Mocco J, Faries P, Merad M, Giannarelli C. 2019. Single-cell immune landscape of human atherosclerotic plaques. Nat Med 25: 1576–88

17. Wirka RC, Wagh D, Paik DT, Pjanic M, Nguyen T, Miller CL, Kundu R, Nagao M, Coller J, Koyano TK, Fong R, Woo YJ, Liu B, Montgomery SB, Wu JC, Zhu K, Chang R, Alamprese M, Tallquist MD, Kim JB, Quertermous T. 2019. Atheroprotective roles of smooth muscle cell phenotypic modulation and the TCF21 disease gene as revealed by single-cell analysis. Nat Med 25: 1280–9

18. Depuydt MA, Prange KH, Slenders L, Ord T, Elbersen D, Boltjes A, de Jager SC, Asselbergs FW, de Borst GJ, Aavik E, Lonnberg T, Lutgens E, Glass CK, den Ruijter HM, Kaikkonen MU, Bot I, Slutter B, van der Laan SW, Yla-Herttuala S, Mokry M, Kuiper J, de Winther MP, Pasterkamp G. 2020. Microanatomy of the Human Atherosclerotic Plaque by Single-Cell Transcriptomics. Circ Res

19. von Scheidt M, Zhao Y, Kurt Z, Pan C, Zeng L, Yang X, Schunkert H, Lusis AJ. 2017. Applications and Limitations of Mouse Models for Understanding Human Atherosclerosis. Cell Metab 25: 248–61

20. Stuart T, Butler A, Hoffman P, Hafemeister C, Papalexi E, Mauck WM, 3rd, Hao Y, Stoeckius M, Smibert P, Satija R. 2019. Comprehensive Integration of Single-Cell Data. Cell 177: 1888–902 e21

21. Korsunsky I, Millard N, Fan J, Slowikowski K, Zhang F, Wei K, Baglaenko Y, Brenner M, Loh PR, Raychaudhuri S. 2019. Fast, sensitive and accurate integration of single-cell data with Harmony. Nat Methods 16: 1289–96

22. Shafer MER. 2019. Cross-Species Analysis of Single-Cell Transcriptomic Data. Front Cell Dev Biol 7: 175

23. Vafadarnejad E, Rizzo G, Krampert L, Arampatzi P, Nugroho VA, Schulz D, Roesch M, Alayrac P, Vilar J, Silvestre J-S, Zernecke A, Saliba A-E, Cochain C. 2019. Time-resolved single-cell transcriptomics uncovers dynamics of cardiac neutrophil diversity in murine myocardial infarction. bioRxiv: 738005

24. Fernandez DM, Rahman AH, Fernandez NF, Chudnovskiy A, Amir ED, Amadori L, Khan NS, Wong CK, Shamailova R, Hill CA, Wang Z, Remark R, Li JR, Pina C, Faries C, Awad AJ, Moss N, Bjorkegren JLM, Kim-Schulze S, Gnjatic S, Ma’ayan A, Mocco J, Faries P, Merad M, Giannarelli C. 2019. Single-cell immune landscape of human atherosclerotic plaques. Nat Med

25. Heberle H, Meirelles GV, da Silva FR, Telles GP, Minghim R. 2015. InteractiVenn: a web-based tool for the analysis of sets through Venn diagrams. BMC Bioinformatics 16: 169

26. Cole JE, Park I, Ahern DJ, Kassiteridi C, Danso Abeam D, Goddard ME, Green P, Maffia P, Monaco C. 2018. Immune cell census in murine atherosclerosis: cytometry by time of flight illuminates vascular myeloid cell diversity. Cardiovasc Res 114: 1360–71

27. Van Hove H, Martens L, Scheyltjens I, De Vlaminck K, Pombo Antunes AR, De Prijck S, Vandamme N, De Schepper S, Van Isterdael G, Scott CL, Aerts J, Berx G, Boeckxstaens GE, Vandenbroucke RE, Vereecke L, Moechars D, Guilliams M, Van Ginderachter JA, Saeys Y, Movahedi K. 2019. A single-cell atlas of mouse brain macrophages reveals unique transcriptional identities shaped by ontogeny and tissue environment. Nat Neurosci 22: 1021–35

28. Rizzo G, Vafadarnejad E, Arampatzi P, Silvestre J-S, Zernecke A, Saliba A-E, Cochain C. 2020. Single-cell transcriptomic profiling maps monocyte/macrophage transitions after myocardial infarction in mice. bioRxiv: 2020.04.14.040451

29. Nugent AA, Lin K, van Lengerich B, Lianoglou S, Przybyla L, Davis SS, Llapashtica C, Wang J, Kim DJ, Xia D, Lucas A, Baskaran S, Haddick PCG, Lenser M, Earr TK, Shi J, Dugas JC, Andreone BJ, Logan T, Solanoy HO, Chen H, Srivastava A, Poda SB, Sanchez PE, Watts RJ, Sandmann T, Astarita G, Lewcock JW, Monroe KM, Di Paolo G. 2020. TREM2 Regulates Microglial Cholesterol Metabolism upon Chronic Phagocytic Challenge. Neuron 105: 837–54 e9

30. Deczkowska A, Weiner A, Amit I. 2020. The Physiology, Pathology, and Potential Therapeutic Applications of the TREM2 Signaling Pathway. Cell 181: 1207–17

31. McArdle S, Buscher K, Ghosheh Y, Pramod AB, Miller J, Winkels H, Wolf D, Ley K. 2019. Migratory and Dancing Macrophage Subsets in Atherosclerotic Lesions. Circ Res 125: 1038–51

32. Weinberger T, Esfandyari D, Messerer D, Percin G, Schleifer C, Thaler R, Liu L, Stremmel C, Schneider V, Vagnozzi RJ, Schwanenkamp J, Fischer M, Busch K, Klapproth K, Ishikawa-Ankerhold H, Klosges L, Titova A, Molkentin JD, Kobayashi Y, Engelhardt S, Massberg S, Waskow C, Perdiguero EG, Schulz C. 2020. Ontogeny of arterial macrophages defines their functions in homeostasis and inflammation. Nat Commun 11: 4549

33. Menezes S, Melandri D, Anselmi G, Perchet T, Loschko J, Dubrot J, Patel R, Gautier EL, Hugues S, Longhi MP, Henry JY, Quezada SA, Lauvau G, Lennon-Dumenil AM, Gutierrez-Martinez E, Bessis A, Gomez-Perdiguero E, Jacome-Galarza CE, Garner H, Geissmann F, Golub R, Nussenzweig MC, Guermonprez P. 2016. The Heterogeneity of Ly6C(hi) Monocytes Controls Their Differentiation into iNOS(+) Macrophages or Monocyte-Derived Dendritic Cells. Immunity 45: 1205–18

34. Bosteels C, Neyt K, Vanheerswynghels M, van Helden MJ, Sichien D, Debeuf N, De Prijck S, Bosteels V, Vandamme N, Martens L, Saeys Y, Louagie E, Lesage M, Williams DL, Tang SC, Mayer JU, Ronchese F, Scott CL, Hammad H, Guilliams M, Lambrecht BN. 2020. Inflammatory Type 2 cDCs Acquire Features of cDC1s and Macrophages to Orchestrate Immunity to Respiratory Virus Infection. Immunity 52: 1039–56 e9

35. Leach SM, Gibbings SL, Tewari AD, Atif SM, Vestal B, Danhorn T, Janssen WJ, Wager TD, Jakubzick CV. 2020. Human and mouse transcriptome profiling identifies cross-species homology in pulmonary and lymph node mononuclear phagocytes. bioRxiv: 2020.04.30.070839

36. Maier B, Leader AM, Chen ST, Tung N, Chang C, LeBerichel J, Chudnovskiy A, Maskey S, Walker L, Finnigan JP, Kirkling ME, Reizis B, Ghosh S, D’Amore NR, Bhardwaj N, Rothlin CV, Wolf A, Flores R, Marron T, Rahman AH, Kenigsberg E, Brown BD, Merad M. 2020. A conserved dendritic-cell regulatory program limits antitumour immunity. Nature 580: 257–62

37. Villani AC, Satija R, Reynolds G, Sarkizova S, Shekhar K, Fletcher J, Griesbeck M, Butler A, Zheng S, Lazo S, Jardine L, Dixon D, Stephenson E, Nilsson E, Grundberg I, McDonald D, Filby A, Li W, De Jager PL, Rozenblatt-Rosen O, Lane AA, Haniffa M, Regev A, Hacohen N. 2017. Single-cell RNA-seq reveals new types of human blood dendritic cells, monocytes, and progenitors. Science 356

38. Dutertre CA, Becht E, Irac SE, Khalilnezhad A, Narang V, Khalilnezhad S, Ng PY, van den Hoogen LL, Leong JY, Lee B, Chevrier M, Zhang XM, Yong PJA, Koh G, Lum J, Howland SW, Mok E, Chen J, Larbi A, Tan HKK, Lim TKH, Karagianni P, Tzioufas AG, Malleret B, Brody J, Albani S, van Roon J, Radstake T, Newell EW, Ginhoux F. 2019. Single-Cell Analysis of Human Mononuclear Phagocytes Reveals Subset-Defining Markers and Identifies Circulating Inflammatory Dendritic Cells. Immunity 51: 573–89 e8

39. Yahagi K, Kolodgie FD, Otsuka F, Finn AV, Davis HR, Joner M, Virmani R. 2016. Pathophysiology of native coronary, vein graft, and in-stent atherosclerosis. Nat Rev Cardiol 13: 79–98

40. Otsuka F, Joner M, Prati F, Virmani R, Narula J. 2014. Clinical classification of plaque morphology in coronary disease. Nat Rev Cardiol 11: 379–89

41. Virmani R, Kolodgie FD, Burke AP, Farb A, Schwartz SM. 2000. Lessons from sudden coronary death: a comprehensive morphological classification scheme for atherosclerotic lesions. Arterioscler Thromb Vasc Biol 20: 1262–75

42. Clement M, Basatemur G, Masters L, Baker L, Bruneval P, Iwawaki T, Kneilling M, Yamasaki S, Goodall J, Mallat Z. 2016. Necrotic Cell Sensor Clec4e Promotes a Proatherogenic Macrophage Phenotype Through Activation of the Unfolded Protein Response. Circulation 134: 1039–51

43. Keren-Shaul H, Spinrad A, Weiner A, Matcovitch-Natan O, Dvir-Szternfeld R, Ulland TK, David E, Baruch K, Lara-Astaiso D, Toth B, Itzkovitz S, Colonna M, Schwartz M, Amit I. 2017. A Unique Microglia Type Associated with Restricting Development of Alzheimer’s Disease. Cell 169: 1276–90 e17

44. Nugent AA, Lin K, van Lengerich B, Lianoglou S, Przybyla L, Davis SS, Llapashtica C, Wang J, Kim DJ, Xia D, Lucas A, Baskaran S, Haddick PCG, Lenser M, Earr TK, Shi J, Dugas JC, Andreone BJ, Logan T, Solanoy HO, Chen H, Srivastava A, Poda SB, Sanchez PE, Watts RJ, Sandmann T, Astarita G, Lewcock JW, Monroe KM, Di Paolo G. 2019. TREM2 Regulates Microglial Cholesterol Metabolism upon Chronic Phagocytic Challenge. Neuron

45. Xiong X, Kuang H, Ansari S, Liu T, Gong J, Wang S, Zhao XY, Ji Y, Li C, Guo L, Zhou L, Chen Z, Leon-Mimila P, Chung MT, Kurabayashi K, Opp J, Campos-Perez F, Villamil-Ramirez H, Canizales-Quinteros S, Lyons R, Lumeng CN, Zhou B, Qi L, Huertas-Vazquez A, Lusis AJ, Xu XZS, Li S, Yu Y, Li JZ, Lin JD. 2019. Landscape of Intercellular Crosstalk in Healthy and NASH Liver Revealed by Single-Cell Secretome Gene Analysis. Mol Cell 75: 644–60 e5

46. Remmerie A, Martens L, Thone T, Castoldi A, Seurinck R, Pavie B, Roels J, Vanneste B, De Prijck S, Vanhockerhout M, Binte Abdul Latib M, Devisscher L, Hoorens A, Bonnardel J, Vandamme N, Kremer A, Borghgraef P, Van Vlierberghe H, Lippens S, Pearce E, Saeys Y, Scott CL. 2020. Osteopontin Expression Identifies a Subset of Recruited Macrophages Distinct from Kupffer Cells in the Fatty Liver. Immunity 53: 641–57 e14

47. Ramachandran P, Dobie R, Wilson-Kanamori JR, Dora EF, Henderson BEP, Luu NT, Portman JR, Matchett KP, Brice M, Marwick JA, Taylor RS, Efremova M, Vento-Tormo R, Carragher NO, Kendall TJ, Fallowfield JA, Harrison EM, Mole DJ, Wigmore SJ, Newsome PN, Weston CJ, Iredale JP, Tacke F, Pollard JW, Ponting CP, Marioni JC, Teichmann SA, Henderson NC. 2019. Resolving the fibrotic niche of human liver cirrhosis at single-cell level. Nature 575: 512–8

48. Jaitin DA, Adlung L, Thaiss CA, Weiner A, Li B, Descamps H, Lundgren P, Bleriot C, Liu Z, Deczkowska A, Keren-Shaul H, David E, Zmora N, Eldar SM, Lubezky N, Shibolet O, Hill DA, Lazar MA, Colonna M, Ginhoux F, Shapiro H, Elinav E, Amit I. 2019. Lipid-Associated Macrophages Control Metabolic Homeostasis in a Trem2- Dependent Manner. Cell 178: 686–98 e14

49. Zhou Y, Song WM, Andhey PS, Swain A, Levy T, Miller KR, Poliani PL, Cominelli M, Grover S, Gilfillan S, Cella M, Ulland TK, Zaitsev K, Miyashita A, Ikeuchi T, Sainouchi M, Kakita A, Bennett DA, Schneider JA, Nichols MR, Beausoleil SA, Ulrich JD, Holtzman DM, Artyomov MN, Colonna M. 2020. Human and mouse single-nucleus transcriptomics reveal TREM2-dependent and TREM2-independent cellular responses in Alzheimer’s disease. Nat Med 26: 131–42

50. Thrupp N, Frigerio CS, Wolfs L, Skene NG, Poovathingal S, Fourne Y, Matthews PM, Theys T, Mancuso R, de Strooper B, Fiers M. 2020. Single nucleus sequencing fails to detect microglial activation in human tissue. bioRxiv: 2020.04.13.035386

51. Krasemann S, Madore C, Cialic R, Baufeld C, Calcagno N, El Fatimy R, Beckers L, O’Loughlin E, Xu Y, Fanek Z, Greco DJ, Smith ST, Tweet G, Humulock Z, Zrzavy T, Conde-Sanroman P, Gacias M, Weng Z, Chen H, Tjon E, Mazaheri F, Hartmann K, Madi A, Ulrich JD, Glatzel M, Worthmann A, Heeren J, Budnik B, Lemere C, Ikezu T, Heppner FL, Litvak V, Holtzman DM, Lassmann H, Weiner HL, Ochando J, Haass C, Butovsky O. 2017. The TREM2-APOE Pathway Drives the Transcriptional Phenotype of Dysfunctional Microglia in Neurodegenerative Diseases. Immunity 47: 566–81 e9

52. Stoeckius M, Zheng S, Houck-Loomis B, Hao S, Yeung BZ, Mauck WM, 3rd, Smibert P, Satija R. 2018. Cell Hashing with barcoded antibodies enables multiplexing and doublet detection for single cell genomics. Genome Biol 19: 224

53. McGinnis CS, Patterson DM, Winkler J, Conrad DN, Hein MY, Srivastava V, Hu JL, Murrow LM, Weissman JS, Werb Z, Chow ED, Gartner ZJ. 2019. MULTI-seq: sample multiplexing for single-cell RNA sequencing using lipid-tagged indices. Nat Methods 16: 619–26

54. Molgora M, Esaulova E, Vermi W, Hou J, Chen Y, Luo J, Brioschi S, Bugatti M, Omodei AS, Ricci B, Fronick C, Panda SK, Takeuchi Y, Gubin MM, Faccio R, Cella M, Gilfillan S, Unanue ER, Artyomov MN, Schreiber RD, Colonna M. 2020. TREM2 Modulation Remodels the Tumor Myeloid Landscape Enhancing Anti-PD-1 Immunotherapy. Cell

55. Katzenelenbogen Y, Sheban F, Yalin A, Yofe I, Svetlichnyy D, Jaitin DA, Bornstein C, Moshe A, Keren-Shaul H, Cohen M, Wang SY, Li B, David E, Salame TM, Weiner A, Amit I. 2020. Coupled scRNA-Seq and Intracellular Protein Activity Reveal an Immunosuppressive Role of TREM2 in Cancer. Cell

56. N AG, Bensinger SJ, Hong C, Beceiro S, Bradley MN, Zelcer N, Deniz J, Ramirez C, Diaz M, Gallardo G, de Galarreta CR, Salazar J, Lopez F, Edwards P, Parks J, Andujar M, Tontonoz P, Castrillo A. 2009. Apoptotic cells promote their own clearance and immune tolerance through activation of the nuclear receptor LXR. Immunity 31: 245–58

57. Liu Z, Gu Y, Chakarov S, Bleriot C, Kwok I, Chen X, Shin A, Huang W, Dress RJ, Dutertre CA, Schlitzer A, Chen J, Ng LG, Wang H, Liu Z, Su B, Ginhoux F. 2019. Fate Mapping via Ms4a3-Expression History Traces Monocyte-Derived Cells. Cell 178: 1509–25 e19

58. Werner Y, Mass E, Ashok Kumar P, Ulas T, Handler K, Horne A, Klee K, Lupp A, Schutz D, Saaber F, Redecker C, Schultze JL, Geissmann F, Stumm R. 2020. Cxcr4 distinguishes HSC-derived monocytes from microglia and reveals monocyte immune responses to experimental stroke. Nat Neurosci 23: 351–62

59. Alsaigh T, Evans D, Frankel D, Torkamani A. 2020. Decoding the transcriptome of atherosclerotic plaque at single-cell resolution. bioRxiv: 2020.03.03.968123

60. Rodriques SG, Stickels RR, Goeva A, Martin CA, Murray E, Vanderburg CR, Welch J, Chen LM, Chen F, Macosko EZ. 2019. Slide-seq: A scalable technology for measuring genome-wide expression at high spatial resolution. Science 363: 1463–7

61. Burke AP, Kolodgie FD, Zieske A, Fowler DR, Weber DK, Varghese PJ, Farb A, Virmani R. 2004. Morphologic findings of coronary atherosclerotic plaques in diabetics: a postmortem study. Arterioscler Thromb Vasc Biol 24: 1266–71

62. Fairweather D. 2014. Sex differences in inflammation during atherosclerosis. Clin Med Insights Cardiol 8: 49–59

63. Wendorff C, Wendorff H, Pelisek J, Tsantilas P, Zimmermann A, Zernecke A, Kuehnl A, Eckstein HH. 2015. Carotid Plaque Morphology Is Significantly Associated With Sex, Age, and History of Neurological Symptoms. Stroke 46: 3213–9

64. Mereu E, Lafzi A, Moutinho C, Ziegenhain C, MacCarthy DJ, Alvarez A, Batlle E, Sagar, Grün D, Lau JK, Boutet SC, Sanada C, Ooi A, Jones RC, Kaihara K, Brampton C, Talaga Y, Sasagawa Y, Tanaka K, Hayashi T, Nikaido I, Fischer C, Sauer S, Trefzer T, Conrad C, Adiconis X, Nguyen LT, Regev A, Levin JZ, Parekh S, Janjic A, Wange LE, Bagnoli JW, Enard W, Gut M, Sandberg R, Gut I, Stegle O, Heyn H. 2019. Benchmarking Single-Cell RNA Sequencing Protocols for Cell Atlas Projects. bioRxiv: 630087

65. van den Brink SC, Sage F, Vertesy A, Spanjaard B, Peterson-Maduro J, Baron CS, Robin C, van Oudenaarden A. 2017. Single-cell sequencing reveals dissociation-induced gene expression in tissue subpopulations. Nat Methods 14: 935–6

